# Drug Repurposing: The Anthelmintics Niclosamide and Nitazoxanide are Potent TMEM16A Antagonists that Fully Bronchodilate Airways

**DOI:** 10.1101/254888

**Authors:** Kent Miner, Katja Labitzke, Benxian Liu, Paul Wang, Kathryn Henckels, Kevin Gaida, Robin Elliott, Jian Jeffrey Chen, Longbin Liu, Anh Leith, Esther Trueblood, Kelly Hensley, Xing-Zhong Xia, Oliver Homann, Brian Bennett, Mike Fiorino, John Whoriskey, Gang Yu, Sabine Escobar, Min Wong, Teresa L. Born, Alison Budelsky, Mike Comeau, Dirk Smith, Jonathan Phillips, James A. Johnston, Joe McGivern, Kerstin Weikl, David Powers, Karl Kunzelmann, Deanna Mohn, Andreas Hochheimer, John K. Sullivan

## Abstract

There is an unmet need in severe asthma where approximately 40% of patients exhibit poor β-agonist responsiveness, suffer daily symptoms and show frequent exacerbations. Antagonists of the Ca^2+^-activated-Cl^−^ channel, TMEM16A, offers a new mechanism to bronchodilate airways and block the multiple contractiles operating in severe disease. To identify TMEM16A antagonists we screened a library of ~580,000 compounds. The anthelmintics niclosamide, nitazoxanide and related compounds were identified as potent TMEM16A antagonists that blocked airway smooth muscle depolarization and contraction. To evaluate whether TMEM16A antagonists resist use- and inflammatory-desensitization pathways limiting β-agonist action, we tested their efficacy under harsh conditions using maximally contracted airways or airways pretreated with a cytokine cocktail. Stunningly, TMEM16A antagonists fully bronchodilated airways, while the β-agonist isoproterenol showed only partial effects. Thus, antagonists of TMEM16A and repositioning of niclosamide and nitazoxanide represent an important additional treatment for patients with severe asthma and COPD that is poorly controlled with existing therapies. It is of note that drug repurposing has also attracted wide interest in niclosamide and nitazoxanide as a new treatment for cancer and infectious disease. For the first time we identify TMEM16A as a molecular target for these drugs and thus provide fresh insights into their mechanism for the treatment of these disorders in addition to respiratory disease.

## INTRODUCTION

Asthma afflicts over 300 million people worldwide, with 5-10% of patients suffering from severe disease which remains poorly controlled despite maximal treatment with an inhaled corticosteroid (ICS) and long-acting β-agonist (LABA). Inflammation is poorly controlled in severe asthmatics, which often necessitates addition of oral corticosteroids, and ~40% of patients show poor β-agonist responsiveness (Moore et al. 2007, Moore et al. 2010). Agonists of the β2-adrenergic receptor (β2AR) signal through the stimulatory G-protein, Gsα, to activate adenylate cyclase (AC), increase cAMP levels and activate PKA. While this mechanism offers the advantage of delivering a negative intracellular signal (cAMP/PKA) to airway smooth muscle (ASM) cells that blocks multiple contractants, repeated β-agonist use and poorly controlled inflammation can cause desensitization of this pathway. Phosphodiesterases (PDE) can degrade cAMP, and increased levels of PDE4D have been detected in biopsies of asthmatic ASM cells leading to reduced β-agonist cAMP responses (Trian et al. 2011). Proinflammatory cytokines or allergic inflammation has been found to increase expression of the inhibitory G-protein, Giα (Hakonarson et al. 1996), uncouple the β2AR from Gsα-induced activation of adenylate cyclase (Shore et al. 1997) and upregulate COX-2 and PGE2 production inducing heterologous β2AR desensitization of ASM cells and reduced β-agonist responsiveness (Moore et al. 2001).

Ion channels represent an alternative target class to modulate contraction which may resist these desensitization pathways, but it has been poorly understood what channels control excitation-contraction coupling in ASM cells. In vascular and airway smooth muscle, bronchoconstrictors have been recognized for years to induce calcium-activated chloride channels (CaCCs) promoting chloride efflux and spontaneous transient inward currents (STICs) contributing to depolarization and contraction, but the molecular identity of the channel has remained elusive (Janssen and Sims 1995, Large and Wang 1996, Wellman and Nelson 2003).

In 2008, three separate groups employing distinct methods identified TMEM16A as the long sought-after calcium-activated chloride channel recognized in secretory epithelial cells, smooth muscle cells and sensory neurons (Caputo et al. 2008, Yang et al. 2008, Schroeder et al. 2008). While these initial studies solidified TMEM16A’s role in the bronchial epithelium as responsible for the elevated Ca^2+^-dependent chloride secretion induced by Th2 cytokines, subsequent studies by Huang et al. (2012) expanded this perspective suggesting TMEM16A may have a dual role in controlling both ASM contraction and epithelial mucin secretion, a hallmark of allergic asthma. Although this research, and more recent findings by Danielsson et al. (2015), introduced TMEM16A antagonists as a novel approach to bronchodilate airways, only a limited number of compounds were explored and details of their relative efficacy compared to β-agonists remained unclear.

To identify additional TMEM16A antagonists and characterize their potential as novel bronchodilators, we screened a library of over half a million compounds. For the first time, we describe the approved drug niclosamide, nitazoxanide and related compounds as potent TMEM16A antagonists that provided robust bronchodilation of airways and resist use- and inflammatory-desensitization pathways limiting β-agonist action. We also discuss separate research from drug-repurposing screens finding the anthelmintics niclosamide and nitazoxanide have efficacy for treating a variety of other disorders, including cancer, and discuss this in the context of TMEM16A, which could represent a molecular target explaining some of these activities.

## MATERIALS AND METHODS

### Materials

CaCCinh-A01, MONNA, and T16Ainh-A01 were purchased from Sigma-Aldrich and tizoxanide was purchased from J&W Pharmlab LLC. Niclosamide (Cpd 1) and niclosamide-related compounds Cpd 2, Cpd 3, Cpd 4, and Cpd 6 - 18, nitazoxanide and CID# 2806957 were from the Amgen small molecule compound collection, as were the benchmark antagonists niflumic acid, dichlorophen, benzbromarone, 1PBC and NTTP. Cpd 5 was synthesized according to a literature procedure (Macielag et al. 1998). The source for other compounds and reagents is provided in each experimental section.

### Cell Line Generation and High-Throughput Screen

The High-Throughput-Screening clone HEK293T:eYFP:TMEM16A(abc) stably expressing eYFP (H149Q,I153L) and human TMEM16A(abc) was generated by stable transfection followed by functional clone selection. HEK293T:eYFP (H149Q,I153L) was cultivated in propagation medium DMEM (PAN #P04-03550) containing 10% FBS (PAN #3302-P281402), 1x Pen/Strep (PAN #P06-07100), 1x L-glutamine (PAN #P04-80100), 10mM HEPES (PAN #P05-01100) and 0.5μg/ml puromycin (Sigma #P7255) and transfected with pIREShyg3:huTMEM16A(abc). Stable cell pool was generated by cultivation in propagation medium containing 0.5μg/ml puromycin and 200μg/ml hygromycin B for 20 days. Cell pool was then diluted in the same medium and seeded in 384-well microtiter plates. Grown clones were replicated into daughter plates using a CyBio Felix pipetting platform (Analytik Jena AG). Daughter plates were analyzed using the eYFP assay essentially as described in the HTS protocol below using a FLIPR-Tetra. Clones showing maximal quenching of eYFP fluorescence intensity after addition of trigger solution (assay buffer containing NaI and ionomycin) were selected for propagation from the mother plate and further characterized with regard to HTS assay development performance parameters including inhibition by the benchmark compound benzbromarone, S/B and relevant HTS quality control statistics (Z-factor, Z’-factor, RZ-factor, RZ’-factor). One clone was finally selected as HTS clone and used throughout the entire HTS campaign to identify TMEM16A antagonists.

For the HTS screen, the screening cell line HEK293T:eYFP(H149Q,I153L):TMEM16A(abc) stably expressing halide sensitive YFP (eYFP (H149Q,I153L)) and human TMEM16A(abc) were propagated in DMEM (PAN #P04-03550) containing 10% FBS (PAN #3302-P281402), 1x Pen/Strep (PAN #P06-07100), 1x L-glutamine (PAN #P04-80100), 10mM HEPES (PAN #P05-01100), 0.5μg/ml puromycin (Sigma #P7255) and 150 μg/ml hygromycin B (Life Technologies #10687-010) in Corning TC-treated flasks. 24 hours prior to the assay 11,000 cells were seeded per well in 30μl assay plating medium (propagation medium containing 0.01% Pluronic F-68 (PAN #P08-2100)) in Corning CellBIND 384-Well Flat Clear Bottom Black Polystyrene Microplates (Corning #3683) and cultivated for 24 hours at 37°C. Before initiating the fully automated and scheduled HTS sequence, cell plates were transferred to an incubator and stored at RT and 0% CO2. Library compounds were delivered in Greiner 384 PP plates (Greiner #784201) containing 1mM compound dissolved in DMSO in columns 1–22 and DMSO in columns 23 and 24 and were stored in a stacker at RT. Further dilutions of compounds were generated by fully automated liquid handling using a CyBio 384-well/25μl pipetting head (Analytik Jena AG) by diluting 0.5μl compound (1mM in DMSO) in 20μl assay buffer (1xHBSS (Life Technologies #14025050) containing 10mM HEPES (PAN #P05-01100) and 0.01% Pluronic F-68 (PAN #P08-2100), adjusted to pH 7.4) in Greiner 384 PP plates (Greiner #784201). Intermediate compound plates contained 25μM library compounds in assay buffer containing 2.5% DMSO in columns 1-22, assay buffer containing 2.5% DMSO as neutral control in column 23 and 100μM Benzbromarone (Sigma #B5774) in assay buffer containing 2.5% DMSO in column 24 as ligand control (benchmark antagonist) and were stored in a stacker at RT. Simultaneous to compound dilution medium was aspirated from cell plates using a Biotek 405 microplate washer followed by washing the cells twice with 65 μl/well assay buffer. With the final aspiration of assay buffer, the residual volume was adjusted to 15μl. 10μl compound solution were then added to the washed cell plates using a CyBio 384-well/25 μl pipetting head (Analytik Jena AG) resulting in a final volume of 25μl assay buffer containing 10μM compound and 1% DMSO. Cell plates were incubated for 30 min at RT and 0% CO2. After incubation cell plates were transferred to a FLIPR-Tetra and 25μl trigger solution (assay buffer containing 20 mM sodium iodide (Sigma #383112) and 4μM ionomycin (Sigma #I0634; stock solution 10mM in DMSO)) was added before reading iodide-quenching of eYFP fluorescence intensity. Final concentration of library compounds in columns 1-22 at read-out was 5μM in the presence of 10mM iodide, 2μM ionomycin and 0.5% DMSO. Neutral control column 23 received no ligand and the ligand control column received a final concentration 20μM benzbromarone in the presence of 10mM iodide, 2μM ionomycin, which corresponded to EC_80_ in this assay setup, and 0.5% DMSO. FLIPR-Tetra settings included excitation wavelength (470-495nm), emission wavelength (515-575nm), 7 reads before dispensing the trigger solution for baseline recording followed by 40 reads after trigger solution dispense. Data analysis was performed using Genedata Screener with aggregation rule “ Max-Min/Max” (Min: baseline from 0 – 10 seconds prior to trigger solution dispense) and normalization based on neutral control (0% inhibition corresponding to 100% TMEM16A activity) and inhibitor control Benzbromarone (100% inhibition corresponding to 0% TMEM16A activity). HTS quality control parameters and statistics (S/B, Z-factor, RZ-factor, Z’-factor and RZ’-factor) were automatically calculated and recorded using Genedata Screener. For unbiased primary hit selection, the POC cut-off (hit threshold) was calculated as Median+3xIQR+20% of all test wells. Confirmation Screening primary was run in triplicates (3 independent, consecutive runs) with confirmed hits defined as median (replicate POC values) > POC cut-off (POC cut-off primary screen). Compounds were selected for dose-response screening after database mining to eliminate frequent hitters and fluorescent primary hit compounds, which were detected by calculating baseline fluorescence intensity measured by FLIPR-Tetra before addition of trigger solution for each compound well. Cut-off was set as Baseline FI <10x STDV of the neutral control. Non-fluorescent compounds were selected and dose response testing was performed with 22-step, 1:2 dilution following the eYFP assay format procedure described above. Threshold was set at IC_50_ <5μM.

The 22 dose response dilutions were also tested in Calcium flux assays using FLIPR Calcium 6 Assay Kit (Molecular Devices # R8190 and R8191) to eliminate compounds, which inhibit ionomycin-dependent increase of intracellular calcium concentration. HEK293T cells were propagated in Corning TC-treated flasks in propagation medium consisting of DMEM (PAN #P04-03550) containing 10% FBS (PAN #3302-P281402), 1x Pen/Strep (PAN #P06-07100), 1x L-glutamine (PAN #P04-80100), 10mM HEPES (PAN #P05-01100), 1x sodium-pyruvate (PAN #P04-43100). Prior to the assay 13,000 cells were plated per well in 30μl propagation medium in Corning Cellbind 384 well microtiter plates (Corning #3683) and cultivated for 24 hours. Cells were stained with labelling dye solution, which was assembled following a protocol provided by manufacturer followed by addition of 8mM probenecid (Sigma-Aldrich #P8761, 250mM stock solution in 1xHBSS (PAN #P04-49505) mixed with an equal volume of 1N NaOH and pH adjusted to 7.4). 25μl labeling dye were transferred per well using Agilent Bravo liquid handling platform followed by incubation for 2 hours at 37°C and 5% CO2. Simultaneously, intermediate compound plates were prepared by transferring 0.64μl of compound solution from compound plates to 15μl assay buffer (1xHBSS (Life Technologies #14025050), 20 mM HEPES (PAN #P05-01100), adjusted to pH 7.4) in columns 1-22 of intermediate plates using a CyBio 384/25μl pipetting head (Analytik Jena AG) followed by storage in a stacker. In order to normalize the assay to identify compounds decreasing calcium signaling final assay concentration of 1μM ionomycin was tested as neutral control in column 23 and buffer only as blank control in column 24. When cell labelling was complete, 10μl of diluted compound were added per well using Agilent Bravo Platform and were incubated for 45 min at room temperature. Cell plates were then transferred to FLIPR-Tetra and 20μl trigger solution (assay buffer containing 4.25μM ionomycin (Sigma #I0634; stock solution 10mM in DMSO) was added to achieve a final Ionomycin concentration of 1μM, which corresponded to the EC_80_ in this assay setup, in the presence of 0.5% DMSO (final). FLIPR-Tetra settings included excitation wavelength (470-495nm), emission wavelength (515-575nm), baseline was recorded before dispensing the trigger solution followed by reading the ionomycin response. Data analysis was performed using Genedata Screener with aggregation rule “ Max-Min/Max” (Min: baseline prior to trigger solution dispense) and normalization based on neutral control (0% inhibition) and blank control without ionomycin (100% inhibition). Dose response cut-off was set at IC_50_<10μM to eliminate compounds, which reduce ionomycin-triggered calcium signaling.

### Cell Line Generation and Medicinal Chemistry Support

The TMEM16A HEK293T eYFP (H149Q, I153L) stable cell line for medchem support was generated by transfecting 10μg of linearized hTMEM16A(abc) DNA constructs into HEK293T eYFP (H149Q,I153L) stable cells. The transfected cells were cultured under 200 μg/ml of hygromycin B selection for 20 days and the stable pool was single cell seeded in 96 well plate and duplicated plate were made after single cell clones were formed. YFP function assay were performed on duplicated plate and clones producing higher changes in fluorescence from the YFP assay were chosen for further expansion to form a stable cell line.

For routine tests of compound activity, HEK293T cells stably co-expressing hTMEM16A (abc) and the halide-sensitive YFP were cultured with DMEM/F-12, HEPES (Life Technologies, Catalog Number: 11330-032) plus 10% (heat inactive) FBS (Life Technologies, Catalog Number: 10082-147), 1X Pen Strep Glutamine (Gibco 10378-016), 0.5μg/ml of puromycin and 200μg/ml Hygromycin B. The day before the assay, 30μl/well of the cell’s suspension was seeded to achieve 15000 cells/well in 384 well plate (Corning^®^ BioCoat^™^ Poly-D-Lysine 384 Well black / clear plate, Cat# 354663). Plates were incubated for 24 hours at 37^0^C with 5% CO_2_. On the assay day, medium was removed and replaced with 20μl/well of assay buffer (Hank’s Balanced Salt Solution with 10mM HEPES pH 7.4), 10μl/well of 40x diluted compounds was added to assay plate and it was incubated at room temperature for 30 min, after which baseline fluorescence was read in the FLIPR instrument for 10 seconds. Then, 5μl/well of a 3x trigger solution (24mM iodide, 3μM ionomycin in assay buffer) was added and the fluorescence kinetic trace was recorded for 2 minutes.

### Q-Patch Electrophysiology for Measuring Compound Effects on TMEM16A

HEK293 cells stably expressing the TMEM16A (acd) variant were purchased from SB Drug Discovery. The HEK293 TMEM16A (abc) stable cell line was from ChanTest and COLO-205 cells were purchased from ATCC. The same standard buffers and recording conditions listed below were used for QPatch studies on all three cell lines.

Recording Solutions: should be made weekly as below and can be stored at room temperature until use. It is recommended not to add ATP to internal solution until immediately before using on the QPatch. The External buffer (in mM) was 140 NaCl, 4 KCl, 2 CaCl2, 1 MgCl2, 10 HEPES, 10 Glucose, pH7.4, while the Internal buffer (in mM) was 110 CsCl, 20 TEA-Cl, 5.374 CaCl2, 1.75 MgCl2, 10 EGTA, 10 HEPES, 4 Na2ATP, pH 7.2.

Once established in whole-cell configuration, cells are clamped to a holding potential (Vhold) of 0 mV on 48 well single hole patch plate. A standardized IV protocol is used to elicit ionic current through the TMEM16A chloride channel at 20 s intervals. Steady-state voltage pulses begin at −100 mV to +100 mV in +20 mV steps for a duration of 500 ms. After each pulse the voltage returns to −100 mV for 50 ms to obtain tail currents, and then returns to the holding potential of 0 mV until the beginning of the next sweep. Block of TMEM16A chloride current due to Test Article is measured at the +100 mV depolarization sweep of the third and final IV protocol run per concentration addition. This ensures a minimum of 60 s per concentration incubation before measuring current block. Currents are acquired at 10 kHz and filtered at 2 kHz, leak subtraction and Rseries are disabled. A 1 MΩ minimum resistance is set because cells are held in an open channel state and therefore resistance is low.

A cumulative concentration response is measured whereby each cell is exposed to five concentrations of test article with a dilution factor of 1:5 (e.g. 0.048, 0.24, 1.2, 6 and 30 μM). Cells are recorded for ~60 s per solution/compound addition. Initially, external solution is applied twice to allow currents to stabilize. Then vehicle (from the saline reservoir) is added twice to monitor any effect 0.3% DMSO might have on currents. This leads directly into the concentration runs by applying Concentration 1 (single addition) for 1 minute. Concentration 2 is then applied for the subsequent minute, etc. Recovery/washout is monitored for a final minute with a final external solution addition.

Peak outward current magnitude at 540ms through 550ms of the IV step is measured for each sweep at 10 s intervals. The final measurement at +100 mV is calculated for each compound application. Each cell’s current magnitude is then normalized to itself at the initial 0.3% vehicle control period prior to compound application. This step is to account for differences in current size for each cell. Normalized responses to test article are plotted against their concentrations to reveal a concentration-inhibition plot.

A Normalized Group Hill fit is then performed on the plotted results to yield a pooled IC_50_ value, reconstructed from a minimum of two cells/concentration. The Baseline Response as well as the Full Response are constrained to 1 and 0, respectively.

### IonWorks Barracuda Electrophysiology Studies

HEK293 cells stably expressing the human TMEM16A (abc) variant and CHO cells expressing the human CFTR gene were from ChanTest. The IonWorks Barracuda (IWB) procedure and data provided are from outsourced studies performed at ChanTest (Charles River Discovery, Cleveland, OH) who were blinded as to the identity of the test articles or compounds provided by Amgen. As a control, benchmark inhibitors CFTRinh-172, GlyH-101 and benzbromarone were included amongst the blinded compounds submitted and performed as expected.

For IonWorks Barracuda studies on TMEM16A compounds, in brief eight test article concentrations were applied to naïve cells (n = 4, where n = the number of replicate wells/concentration) via steel needles of a 384-channel pipettor. Each application consists of addition of 20μl of 2X concentrated test article solution to the total 40μl final volume of the extracellular well of the Population Patch ClampTM (PPC) planar electrode. This addition is followed by mixing (1 times) of the PPC well content. Duration of exposure to each test article concentration was at least five minutes. The electrophysiology procedure used: (a) <u>Intracellular solution</u> containing 50 mM CsCl, 90 mM CsF, 5 mM MgCl2, 1 mM EGTA, 10 mM HEPES, adjusted to pH 7.2 with CsOH; (b) <u>Extracellular solution</u> containing HEPES-Buffered Physiological Saline (HBPS): 137 mM NaCl, 4 mM KCl, 1.8 mM CaCl2, 1 mM MgCl2, 10 mM HEPES, 10 mM glucose, adjusted to pH 7.4 with NaOH; and (c) <u>Ionomycin Stimulation</u> of chloride currents where 10 μM ionomycin is added to all test solutions including vehicle and positive controls. The current was elicited by a 500-ms step pulse to 0 mV followed 1000-ms step pulse to −100 mV from holding potential, −30 mV, with stimulation frequency 0.05 Hz. The specific **Recording Procedure** is as follows: extracellular buffer is loaded into the PPC plate wells (11 μl per well). Cell suspension is then pipetted into the wells (9 μl per well) of the PPC planar electrode. After establishment of a whole-cell configuration via patch perforation, membrane currents are recorded using the on-board patch clamp amplifiers. Recordings (scans) were performed as follows: three scans before and fifteen scans during the five-minute interval after test article application. A full dose-response of benzbromarone was included on each plate as a positive control, while multiple replicates of DMSO were included as negative control. Final DMSO concentration for test and control articles was 0.3%.

For measuring compound effects on CFTR chloride currents, compounds were serially diluted in HEPES-buffered physiological saline to 2X final concentration allowing for an 8-point dose-response analysis. Test article concentrations were applied to naïve cells (n = 4, where n = the number of replicate wells/concentration) via steel needles, where each application will consist of addition of 20μl of 2X concentrated test article solution to a final 40μl volume in the extracellular well of the Population Patch ClampTM (PPC) planar electrode. After mixing (3 times), duration of exposure to compound is at least five minutes. Final solutions contain 0.3% DMSO. The electrophysiology procedure used: (a) <u>Intracellular solution</u> (mM): CsCl, 50; CsF 90; MgCl2, 5; EGTA, 1; HEPES, 10; adjusted to pH 7.2 with KOH, (b) <u>Extracellular, HB-PS Solution</u> (composition in mM): NaCl, 137.0; KCl, 4.0; CaCl2, 1.8; MgCl2, 1; HEPES, 10; adjusted to pH 7.4 with NaOH; and (c) <u>Stimulation</u>, where CFTR current is activated with 20 μM forskolin added to all test solutions including vehicle and positive controls. The current is measured using a pulse pattern consisting of a voltage step to +60 mV, 100 ms duration; voltage ramp from +60 to −120 mV, 1000 ms; step to −120 mV, 200 ms; and step to 0 mV, 200 ms; from holding potential, −30 mV. Current amplitudes were measured at the voltage step to 0 mV. <u>Recording Procedure</u> was as follows: extracellular buffer is loaded into the PPC plate wells (11μl per well). Cell suspension was pipetted into the wells (9μl per well) of the PPC planar electrode. After establishment of a whole-cell configuration via patch perforation, membrane currents were recorded using the patch clamp amplifier in the IonWorks^™^ Barracuda system. Two recordings (scans) were then performed, one scan before and a second scan five minutes after test article application. Every plate included a full dose-response of CFTRinh-172 as a positive control. Data acquisition and analyses were performed using the IonWorks Barracuda^™^ system operation software (version 2.0.2). The decrease in current amplitude after test article application was used to calculate the percent block relative to control. Results for each test article concentration (n ≥ 2) were averaged; the mean and standard deviation values were calculated and used to generate dose-response curves. Inhibition effect was calculated as: % Inhibition = (1 – I_TA_ / I_Baseline_) x 100%, where I_Baseline_ and I_TA_ were the currents measured in control (before addition of a test article) and in the presence of a test article, respectively.

### Histamine-induced Depolarization of Human Bronchial Smooth Muscle Cells

Cultured human bronchial smooth muscle cells (BSMC) from Lonza were resuspended in Smooth Muscle Growth Medium-2 (SmGM-2; Lonza) at a concentration of 4 x 10^5^ cells per ml. One hundred microliters of cells per well were plated in a black wall clear bottom polystyrene 96-well tissue culture plate and incubated in a 37°C humidified incubator with 5% CO2 overnight. Cells were serum starved by removing the SmGM-2 and replaced with one hundred microliters per well of Smooth Muscle Cell Basal Medium 2 (SmBM-2), phenol red-free (PromoCell), and incubated in a 37°C humidified incubator with 5% CO2 for 24 h. SmBM-2 was replaced with one hundred microliters of fresh SmBM-2. Compounds were dissolved in 100% DMSO and serially diluted ½ log in a polypropylene 96-well microtiter plate (drug plate). Columns 6 and 12 were reserved as controls (HI control and LO control respectively) and contained only DMSO. Serially diluted compounds were diluted in SmBM-2 to 10X the final concentration. Twenty-five microliters of 10X compound titrations, and one hundred microliters of Blue dye-loading buffer (FLIPR Membrane Potential Assay Kit; Molecular Devices) were added to the cells. Cells were pre-incubated at room temperature with compound for 0.5 h. Five micromolar (10X) histamine was prepared in SmBM-2. Using the FLIPR-Tetra (Molecular Devices), three measurements of the baseline fluorescence were taken over the span of one minute. Twenty-five microliters per well of 10X histamine were added to the first 11 columns of the plate containing the compound treated cells. Twenty-five microliters of SmBM-2 were added to column 12 for the LO control. Fifty-five fluorescence measurements were taken over the span of fourteen minutes. The area under the curve (AUC) from the fluorescence kinetic traces (normalized to baseline fluorescence) were calculated. The amount of fluorescence in the presence of compound compared with that in the presence of DMSO vehicle alone (HI control) was calculated using the formula: % control (POC) = (compd – average LO)/(average HI – average LO)*100.

While the method above using histamine represents the standard protocol we used to evaluate compound effects in blocking pro-contractile depolarization of human BSMCs, for some experiments provided as supplementary figures we employed the cholinergics methacholine or carbachol instead of histamine, or evaluated if the TMEM16A opener, Eact, could itself act like contractants to induce BSMC membrane depolarization.

### Measuring Bronchodilation of Mouse Tracheal Rings by Wire Myograph

#### Instrument and reagents

A series of three Danish Myograph Technologies (DMT) 620M Multi Wire Myograph Systems instruments were typically used for each experiment, with each instrument containing four chambers and allowing tests on four tracheal rings. The instruments were interfaced with PowerLab 4/35 or PowerLab 8/35 data acquisition systems and a computer running DMT Device Enabler and LabChart Pro v8 (AD Instruments) for automatic recognition of devices and simultaneous recording of data. Carbachol (Cat# C4382) and isoproterenol (Cat# I2760) were from Sigma. Mouse IL-13 and mouse IL-1β were from R&D Systems. Histamine, theophylline and mouse TNFα were from Amgen.

#### Mice

Mice used for wire myograph studies were housed in groups at an AAALAC, International accredited facility. Animals were cared for in accordance with the *Guide for the Care and Use of Laboratory Animals*, 8th Edition. All research protocols were reviewed and approved by the Amgen Institutional Animal Care and Use Committee. Female C57 BL/6 (CRL, >12 weeks of age) were housed in individual ventilated caging (IVC) system on an irradiated corncob bedding (Envigo Teklad 7097). Lighting in animal holding rooms was maintained on 12:12 hr light:dark cycle, and the ambient temperature and humidity range was at 68 to 79°F and 30 to 70%, respectively. Animals had ad libitum access to irradiated pelleted feed^2^ (Envigo Teklad Global Rodent Diet-soy protein free extruded 2020X) and reverse-osmosis (RO) chlorinated (0.3 to 0.5 ppm) water via an automatic watering system. Cages were changed biweekly inside an engineered cage changing station.

#### Standard wire myograph studies for measuring bronchodilation

The trachea from C57 BL/6 mice was dissected and collected in PBS with Mg2+ and Ca2+. After further trimming, two 2mm sections (rings) per trachea were then placed in DMEM media containing PSG/HEPES/AA/NaPyr for 1 hour and used immediately or maintained overnight at 37°C in a humidified incubator with 5% CO2. Rings were then mounted into chambers of DMT wire myograph containing L-shaped mounting pins with 2-3 mN of force applied to secure rings on the pins and the rings were allowed to equilibrate for 25-30 min in a physiological saline buffer (PSS, 130 mM NaCl, 4.7 mM KCl, 1.2 mM MgS04, 14.9 mM NaHCO3, 1.2 mM KH2PO4, 0.026 mM EDTA, 1.6 mM CaCl2, 5.5 mM Dextrose). Throughout the experiment, rings were maintained at physiological temperature and gas conditions by heating buffer reservoir and chamber to 37°C and bubbling a 95% O2, 5% CO2 gas mixture into chamber.

After equilibration, a standard tissue Wake-Up procedure was applied by altering tension and then treatment with elevated potassium as follows: 3 mN tension was applied for 5-10 min, then another 2-3 mN tension to total of 5.5 mN and then to 7-8 mN for another 5 min; the tissue pre-tension was then set to 5 mN and allowed to equilibrate for 5-10 min followed by the removal of the PSS buffer and addition of KPSS (60 mM KCl, 74.7mM NaCl in PSS solution), with force changes monitored until they reached a plateau. The tissue was then washed four times with PSS over 5min and the KPSS treatment and washes repeated; the tissue was then allowed to sit in PSS for 10 min and the tension was set to 5 mN.

To qualify every tissue ring and determine its sensitivity to contractant and reference bronchodilator, 6 ml of PSS was added to each chamber and after 5 min ascending doses of the carbachol (CCh) contractant were added to determine the EC_25_, EC_50_, EC_75_ and EC_95_ for CCh for each ring (see Supplementary Figure 33 for example of raw traces and dose-response), where sufficient time (at least 5 min) was allowed for response to plateau after each dose. At the end of the CCh dose-response study when no further increases in force were observed, the β-agonist and reference bronchodilator isoproterenol was added to 10 μM final concentration to monitor tissue relaxation. The tissue was then washed 3 times and then 3 times more with PSS for 5 min until tension returned to baseline.

Only tissue that passed the qualification tests above advanced to studies on test article compounds. Fresh PSS (6 ml) was then added to each tissue bath, tension was reset to 5 mN and the rings were treated with the calculated EC_25_, EC_50_, EC_75_ or EC_95_ concentration of CCh from GraphPad Prism analysis and allowed to incubate for at least 25 min and a plateau was achieved. For standard ascending dose-response studies on TMEM16A antagonists, an EC_75_ or EC_95_ concentration of CCh was used to pre-contract tracheal rings. Ascending doses of the isoproterenol positive control, the vehicle negative controls or the test compound were then added to appropriate rings and incubated for 10 min or until plateau was achieved after each addition. At the end of the experiment when no further changes were observed to test article or positive control, theophylline was added to 2.2 mM final to fully relax airways. The dose-response curved and EC_50_ values for bronchodilation were derived from GraphPad Prism analysis of the data normalized to theophylline as 100% relaxation.

#### Bronchodilation of tracheal rings as function of differing levels of pre-contraction

Standard procedures as listed above were used to prepare, mount and Wake-Up tracheal rings and the concentrations of carbachol providing EC_25_, EC_50_, EC_75_ or EC_95_ levels of contraction for each ring was determined as described above. Rings with differing levels of CCh pre-contraction were then treated with ascending doses of isoproterenol, benzbromarone, niclosamide or Compound 4 to determine the efficacy of bronchodilation, which was normalized to theophylline added at the end as control of 100% relaxation.

#### Effects of cytokines on compound efficacy in relaxing mouse tracheal rings

Airway rings were treated overnight in 2 ml DMEM + PSG/HEPES/AA/NaPyr buffer alone or with the same solution supplemented with 100 ng/ml of mTNFα, mIL-13 and mIL-1β. Unlike like the typical Wake up protocol and ring qualifying tests, mouse tracheal rings were not exposed to KPSS and rings were not observed to relax with a β-agonist following discovery of CCh dose-responses. After several washes with PSS, EC_75_ concentrations of CCh were used to pre-contract rings for >30min alone or in the presence of cytokines. Ascending doses of isoproterenol, benzbromarone, or niclosamide were given to determine efficacy of bronchodilation normalized to theophylline as control of 100% relaxation.

#### Evaluating β-agonist short-term use-dependent desensitization

Standard procedures were used to prepare and mount tissues. However, similar to the experiments assessing the effects of cytokines in relaxing tracheal rings, no KPSS or qualifying tests with isoproterenol to observe relaxation following discovery of CCh dose-responses. To assess use-dependent desensitization, EC_50_ concentrations of CCh were used to pre-contract rings followed by a dose-response with isoproterenol. Rings were then washed several times with PSS, pre-contracted again with EC_50_ CCh, and given another round of isoproterenol dose-response normalized to theophylline as control of 100% relaxation.

#### Assessing compound duration of action in bronchodilating

Mouse airway rings were exposed to a typical Wake up procedure including the use of KPSS and discovery of EC_75_ CCh dose responses. However, no isoproterenol was used to observe tissue relaxation as part of the qualification procedure. With pre-contraction with EC_75_ CCh for >30min, rings were subjected to a compound dose-response, followed by a complete washout of compound, and re-contracted with EC_75_ CCh. Traces were recorded for several hours thereafter and normalized to theophylline as control of 100% relaxation.

### Measuring Bronchodilation of Human Bronchial Rings by Wire Myograph

#### Outsourced bronchodilator studies performed at Biopta (Glasgow, United Kingdom)

Details of bronchodilator studies shown in **Figures 5H,6** and Supplementary Figure 40 are provided below as part of the end assay report on Study CUR011 by Lee Christie (Biopta). Donors with any of the following conditions were excluded from the study: asthma, COPD, emphysema, lung cancer, Cystic Fibrosis, pulmonary fibrosis, pulmonary hypertension, pneumonia. Donors were also excluded if they had smoked in the past 12 months. Donors were also excluded if they had smoked in the past 12 months. Any macroscopically diseased/necrotic tissue was rejected. Furthermore, any tissues that did not respond to functional checks were rejected.

Human airway rings were set up under isometric conditions on a wire myograph in order to examine the influence of the test articles on bronchodilation. In order to assess tissue viability, the airways were challenged with carbachol (10 μM) and then isoprenaline (10 μM) to assess their constriction and relaxation responses, respectively. Airways that did not respond to these initial checks were not used.

Viable airways were pre-constricted with histamine (10 μM), then exposed to one of the following cumulative concentration response curves (CCRCs): positive control, vehicle (water, volume matched to positive control), test article, vehicle (DMSO concentration matched to test articles). At the end of each CCRC, theophylline (2.2 mM) was added to induce maximal relaxation of the tissue.

Specific methodology was as follows: (**a**) Quaternary branches of human airway rings were dissected free from surrounding parenchyma, cut into 2 mm rings and mounted in 5 mL organ baths containing physiological saline solution (composition: 119.0 mM NaCl, 4.7 mM KCl, 1.2 mM MgS04, 24.9 mM NaHCO3, 1.2 mM KH2PO4, 2.5 mM CaCl2, 11.1 mM glucose and 5 μM indomethacin), aerated with 95% O2/5% CO2, and warmed to 37°C. The tissues were allowed to equilibrate for approximately 30 minutes, with washes approximately every 10 minutes, before being set to a tension of approximately 1.0 - 1.5 g and then allowed to equilibrate for a further 90 minutes with washes approximately every 15 minutes. Airways were re-tensioned to 1.0 – 1.5 g if the tension had dropped below 1.0 g during the first 30 minutes of equilibration. (**b**) Airways were exposed to carbachol (bath concentration 10 μM) in order to measure their contractile responses and then isoprenaline (10 μM) in order to assess their relaxation. Airways were washed out and allowed to return to baseline. (**c**) Airways were pre-constricted with histamine (10 μM) before conducting a CCRC to the test article, the positive control (isoprenaline), test article vehicle or positive control vehicle. It should be noted there were no significant effect by any of the vehicles in the studies reported.

#### Bronchodilator studies performed internally at Amgen (Thousand Oaks, CA, USA)

Bronchodilator studies using human bronchial rings (**Figures 5E-5G**, Supplementary Figure 38) used the same instruments and reagents described earlier for studies on mouse tracheal rings.

Non-transplantable human lungs were obtained through IIAM (Edison, NJ) from non-smoking donors who had been ventilated for <3 days with acceptable blood gases and used within 24 hours of cross-clamp time.

Human 4th order bronchial rings were isolated, sliced into 2mm long rings and then mounted individually in chambers of wire myographs containing 6 ml room temperature PSS per chamber. Chambers are aerated with 95% O_2_/5% CO_2_ throughout the experiment. Bronchial rings are allowed to equilibrate in PSS while chambers are warmed to 37°C. Tension on the airway rings was gradually increased until reaching a steady state passive tension of 9.8 mN. The rings were then equilibrated for additional 40-60 minutes, changing buffer every 15-20 minutes. Tension was adjusted if it dropped below 9.8 mN. Airway rings were “ woken up” by exposing to pre-warmed, aerated KPSS and allowing rings to reach plauteau of constriction, followed by washing rings 4 times with PSS. This procedure was repeated 2 additional times.

Airway contraction response was then assessed by treating with increasing doses of the contractant, carbachol (10 nM to 10 μM), waiting for response plateaus (at least 5 minutes between additions). Airways are then relaxed by addition of 10 μM isoproterenol. Airways that demonstrated expected contraction and relaxation were used to evaluate test compounds after a washout and re-equilibration period.

Airway rings were contracted with EC_75_ of carbachol and then exposed to one of the following CCRCs: vehicle (DMSO concentration matched to test article), isoproterenol (positive control), test compound. At the end of each CCRC, rings were fully relaxed by addition of 2.2 mM theophylline. One hundred percent contraction was calculated by subtracting the stable tension remaining in the airway ring after theophylline addition from the tension achieved after addition of EC_75_ carbachol while 100% relaxation is the net tension remaining after theophylline addition. Dose response curves and EC_50_ values for bronchodilation were determined in GraphPad Prism.

### Evaluating the TMEM16A expression in airway smooth muscle cells and the bronchial epithelium

#### RNA sequencing

The RNA-Seq Amgen lung cell dataset containing primary human bronchial epithelial cells and airway smooth muscle cells has been described earlier, as has the method for RNA sequencing (Aisenberg et al. 2016), with modifications and additional details provided below. Similar RNA sequencing and data analysis methods were applied to generate the new RNA-Seq dataset described here for untreated or IL-13 treated mature bronchial epithelial ALI cultures.

Data were analyzed using the Array Studio (Omicsoft, NC) platform, as previously described (Aisenberg et al. 2016), but using v9.0 of the Oshell software and the GENCODE (Harrow et al. 2012) gene model (release 24; Comprehensive version was used for alignment, and Basic version was used for quantification). FPKM values were normalized using a modified version of upper-quartile normalization (Mortazavi et al. 2008, Robinson and Oshlack 2010) in which the 70^th^ percentile FPKM among genes was fixed at a value of 10 (excluding those genes belonging to families with high homology or with maximal transcript length <500 bp). Data from the Cancer Cell Line Encyclopedia (Barretina et al. 2012) were processed by OmicSoft using the same analysis pipeline and normalized using the same methods.

#### Generating human bronchial epithelial ALI cultures and effects of Th2 cytokines

To determine the effects of IL-13 on TMEM16A versus Muc5AC mRNA expression over time in the human bronchial epithelium and evaluate TMEM16A alternative splicing, normal and COPD human bronchial epithelial cells from Lonza were grown on 6.5mm permeable supports (Corning Transwell 3470) submerged in apical and basolateral growth media (BEGM, Lonza, CC-3170), until confluent at ~5 days. Once confluent, apical media was removed, and basolateral growth media was replaced with ALI maintenance media (PneumaCult, STEMCELL Technologies #05001) to initiate the air-liquid interface (ALI). Cultures were then maintained at ALI for ~21 days, with basolateral maintenance media being replenished on Monday-Wednesday-Friday schedule, to generate fully differentiated human bronchial epithelial cells. Cultures were then left untreated or treated with 20 ng/ml IL-13 for 1, 3, 5 or 7 days. For each time point, RNA was prepared from the ALI cultures for the NextGen RNA sequencing following the Qiagen RNeasy Mini kit (cat # 74104) protocol. Each ALI culture was solubilized with 350 μl of Qiagen lysis buffer RLT on the apical transwell and immediately spun through a Qiashredder (Qiagen # 79654) at 15,000 rpm for 2 minutes. The samples were either stored at −80°C and processed at a later date or immediately prepared following the Qiagen RNeasy Mini kit protocol including Part 2 which contains the Qiagen RNase-Free DNase set (Cat # 79254) treatment. Total RNA was eluted in 45 μl of RNase free water supplied from the kit and quantified on a Nanodrop ND-1000 spectrophotometer.

TMEM16A protein expression was determined by immunohistochemistry of untreated or IL-4 or IL-13 treated human bronchial epithelial ALI cultures. Fully differentiated human bronchial epithelial ALI cultures were generated using methods similar to above, except cultures were expanded on permeable supports for 3 days and maintained at ALI for 31 days prior to treatment. At 31 days post-airlift, cultures were left untreated or treated with either 20 ng/ml IL-4 or 20 ng/ml IL-13 for 1, 2, 3, or 5 days. For each timepoint, Transwells were fixed both apically and basolaterally with 2% paraformaldehyde for 2 hours at room temperature, then washed with 70% ethanol. Permeable membranes containing the ALI cultures were then cut away from the plastic insert using a scalpel and processed for immunohistochemistry as described below.

#### TMEM16A expression in tissue isolated from naïve or asthmatic cynomolgus monkeys

Tissue from naïve or asthmatic cynomolgus monkeys was obtained from Charles River Laboratories. Naïve adult cynos were never challenged with *Ascaris suum* aerosol, were from the standard CRL colony and were negative by intradermal screening for *A. suum*. Adult cynos exhibiting sensitivity to *A. suum* antigen and characterized as asthmatic, presumably due to early exposure and allergy to A. *suum* or similar parasite, were maintained in a separate colony and confirmed over time as reproducible sensitivity to an *A. suum* aerosol challenge. Naïve, unchallenged asthmatic (>2 weeks), asthmatic acute challenged (4 hours post aerosol A. suum) and asthmatic subacute challenged (24 hours post *A. suum* aerosol) cynos were euthanized and lung vasculature infused with cold HypoThermosol biopreservation media. Lungs and trachea were shipped on ice packs overnight and necropsy was immediately performed on arrival to prepare specimens for immunohistochemistry and bronchodilation studies.

#### Immunohistochemistry

The routinely formalin-fixed, paraffin embedded tissue blocks were sectioned at a 4 μm thickness and processed for IHC. Paraffin was removed from the tissue sections with xylene and the sections were rehydrated with graded ethanol and immersed in distilled water. Antigen retrieval was performed using Diva Decloaker pretreatment reagent (Biocare Medical, Concord, CA) in a Biocare Decloaking Chamber (Biocare Medical, Concord, CA) set to reach 125°C for 30 seconds, then 90°C for 10 seconds. Tissue sections were processed at room temperature in a Lab Vision 720 automated staining instrument (Thermo Scientific, Waltham, MA). Endogenous peroxidase was blocked using Hydrogen Peroxide Block (cat# TA-125-HP, Thermo Scientific, Waltham, MA) for 10 minutes. Protein Block (cat# X0909, Dako North America) was applied to the tissue sections for 10 minutes. Tissue sections were incubated with anti-human TMEM16A (cat# Ab53212, Abcam, San Francisco, CA). Envision+ System HRP labelled Polymer (cat# K4003, Dako North America) was used to detect the primary antibody. Sections were incubated with DAB plus chromogen substrate (cat#K3468) for 5 minutes.

### Curve Fitting, Statistical Analysis and Small Molecule Informatics

Data were analyzed using GraphPad Prism v7.0 (GraphPad Software, San Diego, CA, USA). All data is expressed as mean ± SEM, unless otherwise indicated. Concentration-response curves were fit by non-linear regression. Data were compared using the Student’s t-test for unpaired samples or by two-way ANOVA using Dunnett’s or Sidak’s multiple comparison tests, as appropriate. P values < 0.05 were considered statistically significant.

Compound lipophilicity was determined using the SwissADME free web tool (Daina, Michielin, and Zoete 2017) and the XLogP3 additive model for logP calculation (Cheng et al. 2007).

## RESULTS

### High throughput screening of small molecule library identifies niclosamide as a novel TMEM16A antagonist

To identify novel inhibitors of the calcium-activated chloride channel TMEM16A, we generated a stable cell line HEK293T:eYFP:TMEM16A(abc) co-expressing the ‘abc’ splice variant of human TMEM16A along with the halide-sensitive YFP mutant [H148Q,I152L] referred to as eYFP. To this end, HEK293T:eYFP cells, which show no significant response to ionomycin and iodide, were stably transfected with a TMEM16A(abc) expression cassette. Clones suitable for high-throughput screening were identified by functional clone selection using the eYFP assay (data not shown). For the HTS eYFP assay, cells were incubated with the test compounds for 30 min and then treated with 10 mM iodide and 2 μM ionomycin. Ionomycin triggers the Ca^2+^-dependent activation of TMEM16A, which allows iodide ions to enter the cell and quench eYFP fluorescence. eYFP assay results were normalized based on the benchmark TMEM16A inhibitor benzbromarone providing 100 Percent of Inhibition (POI) of TMEM16A activity. This corresponds to 0 Percent of Control (POC) response of Ca^2+^-dependent TMEM16A activity.

A library of ~580,000 compounds was screened at a 5 μM final concentration in the TMEM16A halide-sensitive YFP assay. A summary of the screen and the hit triaging process is provided in Supplementary Table 1 and Supplementary Figure 1, and described further below. A total of 1,445 primary hits were identified using a hit cut-off of POC<72.6%, which reflects compounds providing >27.4% inhibition. All primary hits were tested again in three independent, consecutive runs and confirmed 673 hits. In order to eliminate false-positive TMEM16A inhibitors, auto-fluorescent compounds were removed revealing 491 non-fluorescent hits. A Ca^2+^ flux counterscreen was run to eliminate compounds non-specifically interfering with ionomycin-triggered calcium signaling further reducing the number of hits to 328 compounds. Dose response testing of these hits in the TMEM16A eYFP assay yielded 145 hits with an IC_50_<5μM and finally 130 hits that passed quality control by mass spec analysis. **Figure 1A** shows the distribution of hit potency versus average POC response in the HTS assay. Niclosamide (Compound 1) and a related analog, Compound 4, were identified for the first time from our screen as potent TMEM16A inhibitors. Niclosamide with an IC_50_ of 140 nM in the eYFP HTS assay (**Figure 1A**) represents one of the most potent TMEM16A antagonists described to date. Compound 4, a structurally related analog (**Figure 2**), showed lesser activity with an IC_50_ of 970 nM, but both this compound and niclosamide showed only partial block of the halide-sensitive YFP response with a POC of 61.1 ± 7.8 and 69.0 ± 3.3, respectively, reflecting just 38.9% and 31.0 % inhibition. In fact, many of the hits from the screen provided less than 50% inhibition of the iodide eYFP quenching after ionomycin-activation of TMEM16A (**Figure 1A**). This contrasts with the benchmark antagonist benzbromarone, which fully blocked the halide-sensitive response and thus served as a positive control for 100% inhibition (POC = 0). Interestingly, the TMEM16A inhibitor 1PBC was also included in our small molecule library and showed an activity of 74.8 POC (**Figure 1A**) corresponding to 25.2% inhibition. To characterize this compound further we performed 22-point dose-response analysis and found that 1PBC inhibited TMEM16A with IC_50_ of 1.05 μM but only partially blocked the halide-sensitive response just like niclosamide.

**Table 1.**
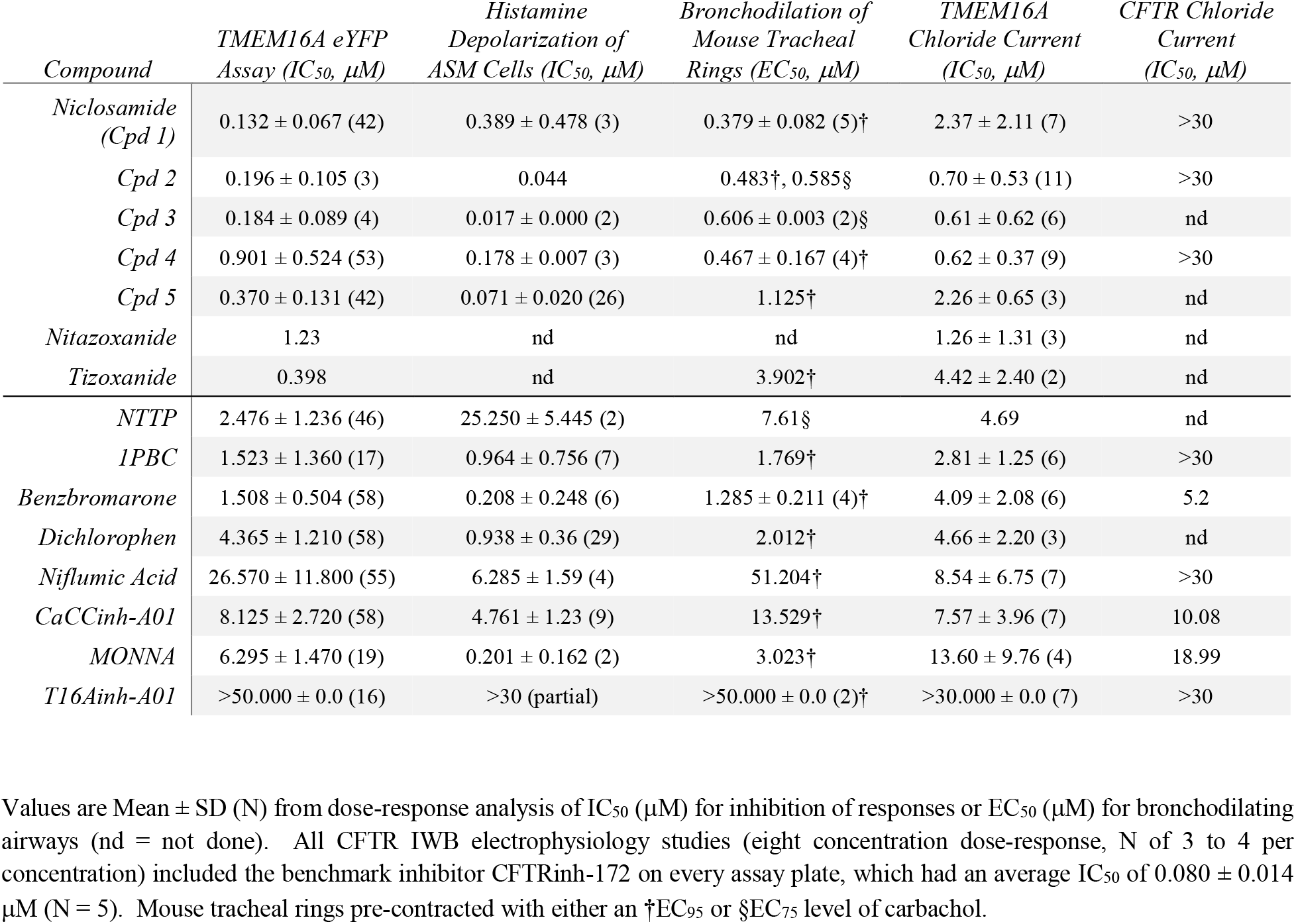
Niclosamide and related compounds are potent inhibitors of TMEM16A conductivity that block histamine depolarization of human airway smooth muscle cells and relax mouse tracheal rings pre-contracted with carbachol.

**Figure 1.**
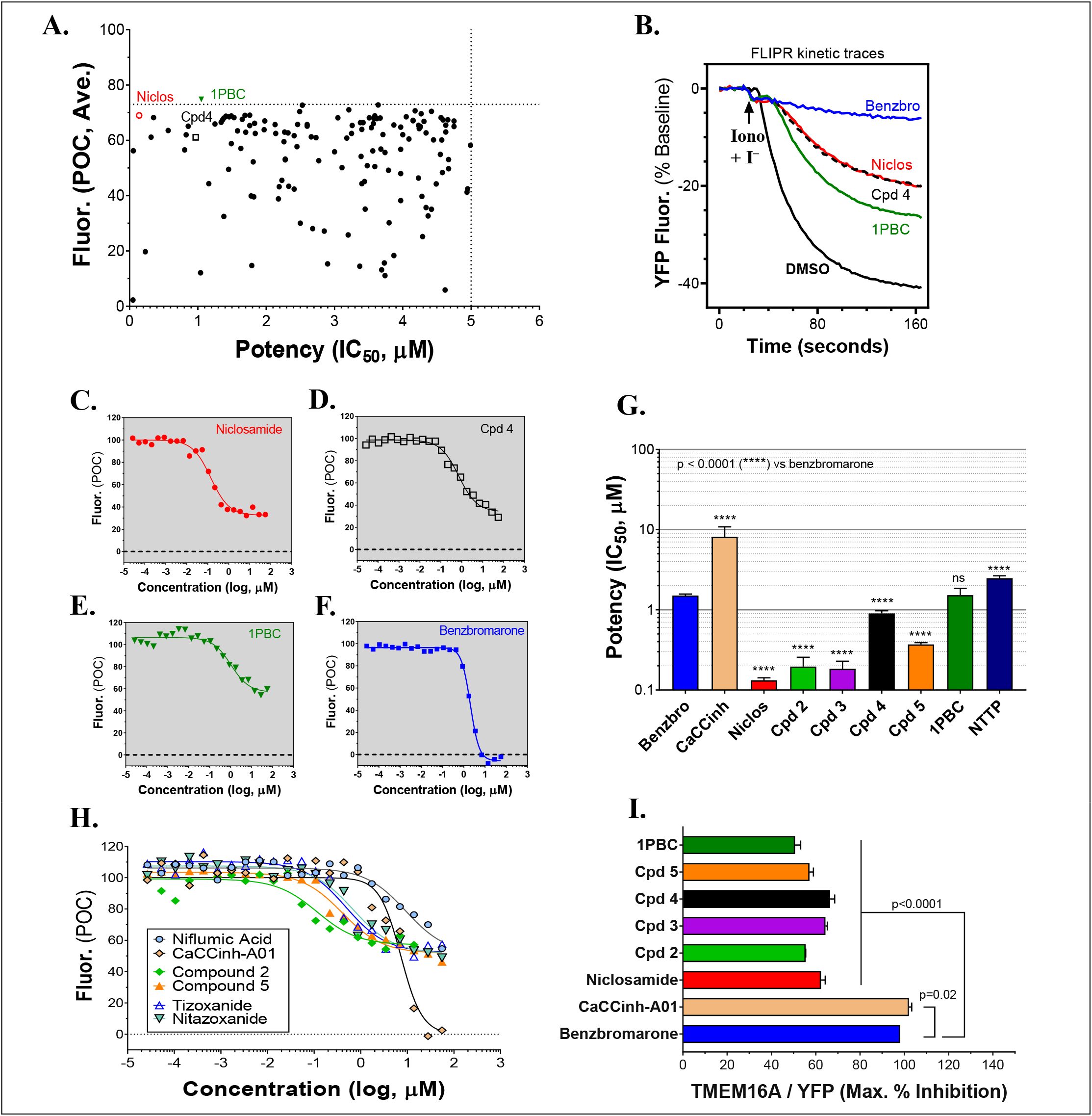
The hierarchical DDM. High-throughput screen using a halide-sensitive YFP assay identifies niclosamide as a highly potent, nanomolar inhibitor of TMEM16A. (**A**) Potency range of the 145 hits from the TMEM16A high-throughput screen and triplicate hit confirmation at 5 μM (n of 4) that reduced the YFP quenching by less than 72.6 percent of control (POC) response to ionomycin (Iono)/iodide alone. The dotted horizontal and vertical lines demarcate the cut-off values (POC < 72.6, IC_50_ < 5 μM) for validated hits. Representative data from the medchem TMEM16A YFP assay showing FLIPR kinetic traces for the DMSO vehicle control or compounds at 55.6 μM (**B**), and dose-response curves indicating niclosamide (**C**), Cpd 4 (**D**) and 1PBC (**E**) only partially blocked the ionomycin-induced YFP response, while benzbromarone (**F**) and CaCCinh-A01 (**H**) gave full block. Niclosamide and related compounds, Cpd 2 - Cpd 5, were more potent TMEM16A antagonists than benchmarks (G,H), but only partially blocked the iodide/YFP response (**I**). Nitazoxanide and tizoxanide were identified as additional inhibitors of TMEM16A (**H**). Mean ± SEM, n of 3-58. **** P < 0.0001; significant difference from the benzbromarone benchmark antagonist; ns, not significant (unpaired *t*-test).

**Figure 2.**
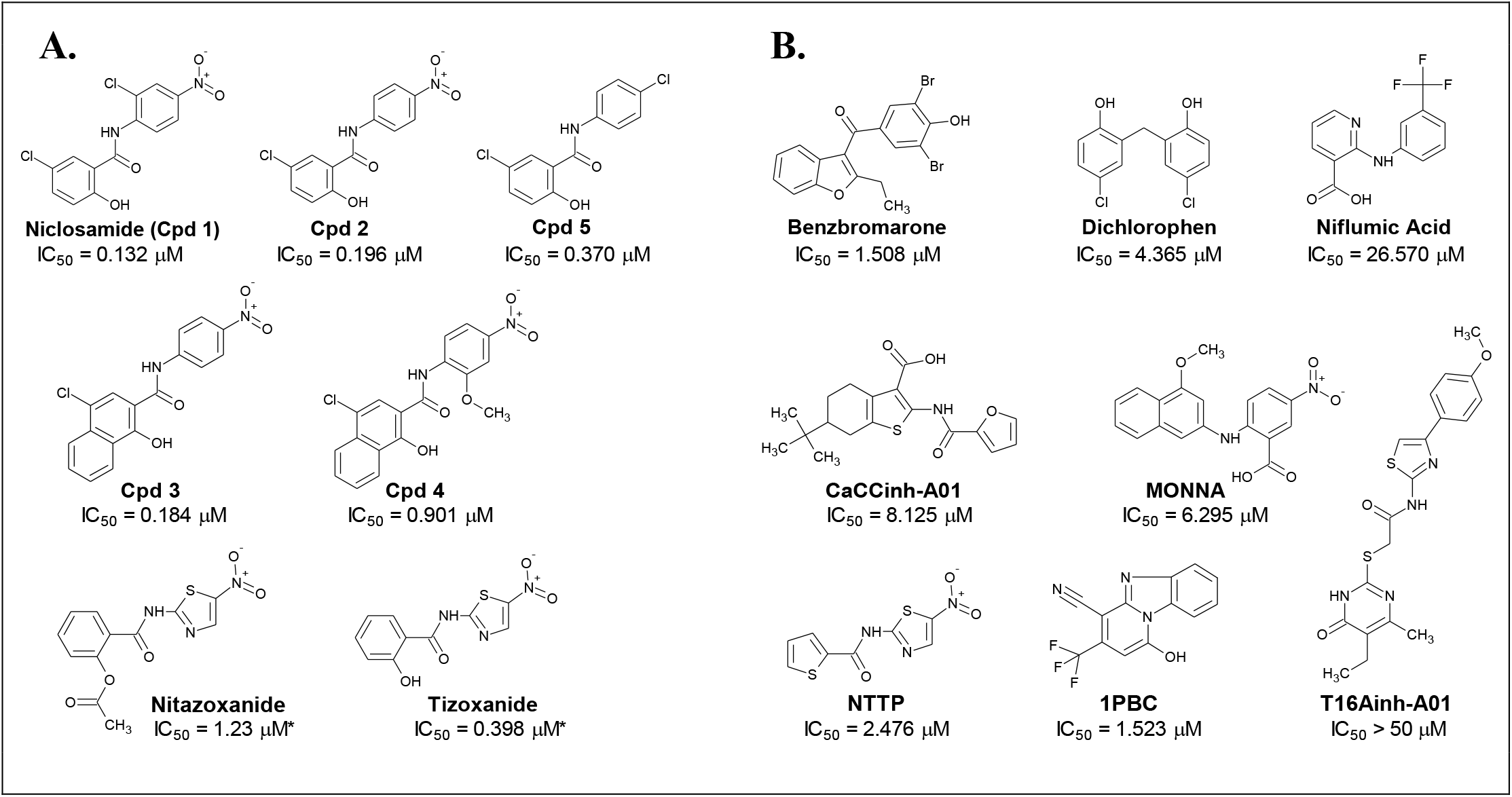
Structures of small molecule antagonists and their activity in the TMEM16A halide-sensitive YFP assay. Molecular structure of the hits niclosamide and Cpd 4 identified from our high-throughput screen and related compounds discovered during our hit-to-lead process, which includes the approved drug nitazoxanide and its metabolic product tizoxanide (**A**). The structures of TMEM16A benchmark antagonists described earlier are shown in panel (**B**) on the right. The average potency of each compound in inhibiting the TMEM16A halide-sensitive YFP response was determined by 22-point dose-response studies, with the average IC_50_ (μM) listed beneath each compound (n = 3-58; with exception of compounds labeled with * which were single determinations).

Side by side comparison of TMEM16A antagonists using HEK293T:eYFP:TMEM16A(abc) cells in medchem follow-up assays revealed that treatment with 1PBC, Cpd 4 and niclosamide caused only partial inhibition of TMEM16A activity whereas benzbromarone blocks almost completely (**Figure 1B**). This was further corroborated in 22-point dose-response analyses showing that the benchmark antagonist benzbromarone fully inhibited the iodide eYFP quenching after ionomycin-activation of TMEM16A (**Figure 1F**), while niclosamide, Cpd 4 and 1PBC showed only partial inhibition even at the highest concentration of 55.6 μM (**Figure 1C-E**). The scope of their block in this follow-up characterization was more pronounced compared to inhibition during HTS. For instance, niclosamide provided about 31% inhibition during HTS but achieved up to 67% inhibition in the dose-response analysis (**Figure 1C**). However, its potency was unchanged. Remarkably, niclosamide with an average IC_50_ of 132 ± 67 nM (n=42) was strikingly more potent than the benchmark antagonist benzbromarone having an average IC_50_ of 1.508 ± 0.504 (n=58) μM (**Figure 1G**, Supplementary Fig. 2). Compound 4 revealed an average IC_50_ of 0.901 ± 0.524 μM (n=53) in comparison to 1PBC with an average IC_50_ of 1.523 ± 1.360 μM (n=17) and CaCCinh-A01 with an average IC_50_ of 8.125 ± 2.720 μM (n=17).

### Activity of additional niclosamide-related compounds and identification of the approved drug nitazoxanide as a new TMEM16A antagonist

The structure and activity of additional niclosamide-related compounds are provided in **Figure 2A**. Compounds 2, 3 and 5 provided sub-μM block of the TMEM16A eYFP response and improved activity compared to benzbromarone (**Figure 1G**), but like niclosamide only partially inhibited the ionomycin-induced eYFP quenching by *iodide* (**Figures 1H,I**). Besides benzbromarone, we also evaluated the TMEM16A antagonists dichlorophen, CaCCinh-A01, MONNA, niflumic acid and T16Ainh-A01. The TMEM16A antagonist CaCCinh-A01 fully inhibited the ionomycin-induced iodide eYFP quenching (**Figures 1H,I**) like benzbromarone, as did the benchmark antagonists MONNA and dichlorophen which had an average IC_50_ of 6.295 ± 1.470 (n=19) and 4.365 ± 1.210 μM (n=58), respectively, in the halide-sensitive YFP assay (**Figure 2**, Supplementary Figure 3). Niflumic acid showed lesser activity with an average IC_50_ of 26.570 ± 11.800 μM (n=55) while the compound T16Ainh-A01 was inactive despite repeat tests, IC_50_ >50 μM (n=16) (Supplementary Figure 3).

The niclosamide-related Compounds 2-5 represent only a small subset of compounds we’ve tested in this series to develop a structure-activity relationship (Supplementary Figure 4). Compounds 2 and 5 are instructive in showing the nitro group is unnecessary for TMEM16A bioactivity and can be replaced with a chlorine atom (**Figure 2A**). The World Health Organization includes niclosamide on its list of Essential Medicines based on its efficacy, safety and cost-effectiveness. Another approved drug, nitazoxanide, is synthesized using the scaffold of niclosamide. Since nitazoxanide and its metabolic product tizoxanide appear structurally similar to niclosamide (**Figure 2A**), we tested these drugs for activity in blocking TMEM16A. Both nitazoxanide and tizoxanide were found to be antagonists of TMEM16A (**Figure 1H**) and exhibit a pharmacology similar to niclosamide in partially inhibiting the iodide/eYFP response.

The TMEM16A inhibitor, NTTP, is noteworthy as it shares the mechanism of 1PBC in blocking TMEM16A by binding four basic residues in the TMEM16A selectivity filter. We have confirmed NTTP is a TMEM16A antagonist (Supplementary Figure 3). Interestingly, an examination of the chemical structure of NTTP reveals it is a highly similar analog of tizoxanide (**Figure 2**). This suggests nitazoxanide and its metabolic product tizoxanide may share the mechanism of NTTP in binding the pore region of TMEM16A to block channel conductivity.

In summary, the direct comparison of niclosamide with eight distinct TMEM16A benchmark antagonists revealed that niclosamide with an IC_50_ of 132 nM in the halide-sensitive YFP assay is 10 to 200 times more potent than other antagonists. Our results further suggest niclosamide and related compounds may utilize a unique mechanism for channel block. Therefore, we performed additional experiments to characterize their impact on chloride currents and effects in blocking airway smooth muscle cell depolarization and contraction of airways.

### Analysis of niclosamide, nitazoxanide and related compounds for inhibition of TMEM16A calcium-activated chloride currents

While our screen identified potent TMEM16A antagonists, we wondered why many just partially inhibited the ionomycin-induced quenching of the eYFP halide sensor by *iodide* in the YFP assay (**Figures 1H,I**). Therefore, we asked how these TMEM16A antagonists perform when measuring their efficacy in blocking calcium-activated *chloride* currents. Electrophysiology studies on this channel, however, can be notoriously difficult as TMEM16A exhibits rapid calcium-dependent inactivation (Wang and Kotlikoff 1997, Tian, Schreiber, and Kunzelmann 2012). An additional challenge is that the ephys assay must have enough throughput to test numerous compounds and quickly return data to the chemist to guide compound synthesis and the hit-to-lead (HTL) process. Because no such assay was yet available, we developed an automated QPatch electrophysiology assay that enables prolonged recordings and potency determinations on up to 100 compounds per week. Whole-cell currents from HEK293 cells stably-transfected with TMEM16A (acd) were recorded using 170 nM free intracellular calcium and 20 mV steps from −100 to +100 mV from a holding potential of 0 mV (**Figure 3A**). Three voltage protocols were run per sample addition allowing at least 60 s incubation time following each vehicle or compound addition. The currents were voltage-dependent and outward rectifying as expected for TMEM16A recordings in low intracellular calcium. Importantly, the currents were stable for about 10 minutes and insensitive to the DMSO vehicle control as shown in the representative QPatch instrument recording in **Figure 3B** where the dots reflect the measured current after each 20 mV step from −100 to +100 mV and three IV protocols are shown for each sample addition. While the optimized assay typically showed <20% rundown over this time frame, it should be noted significant rundown becomes an issue with recordings that are longer than 12-15 minutes. The benchmark TMEM16A antagonist benzbromarone fully inhibited both the outward and inward currents (**Figures 3C,D**). Typical QPatch recording for dose-response analysis included two saline and DMSO additions to validate current stability and voltage-dependence, followed by five concentrations of antagonists as shown in **Figure 3C**. The sustained TMEM16A current following the second DMSO addition and third IV protocol served as the baseline current for calculating percent inhibition. Routine calculations of compound potency measured their effect in reducing the outward current at +100 mV.

**Figure 3.**
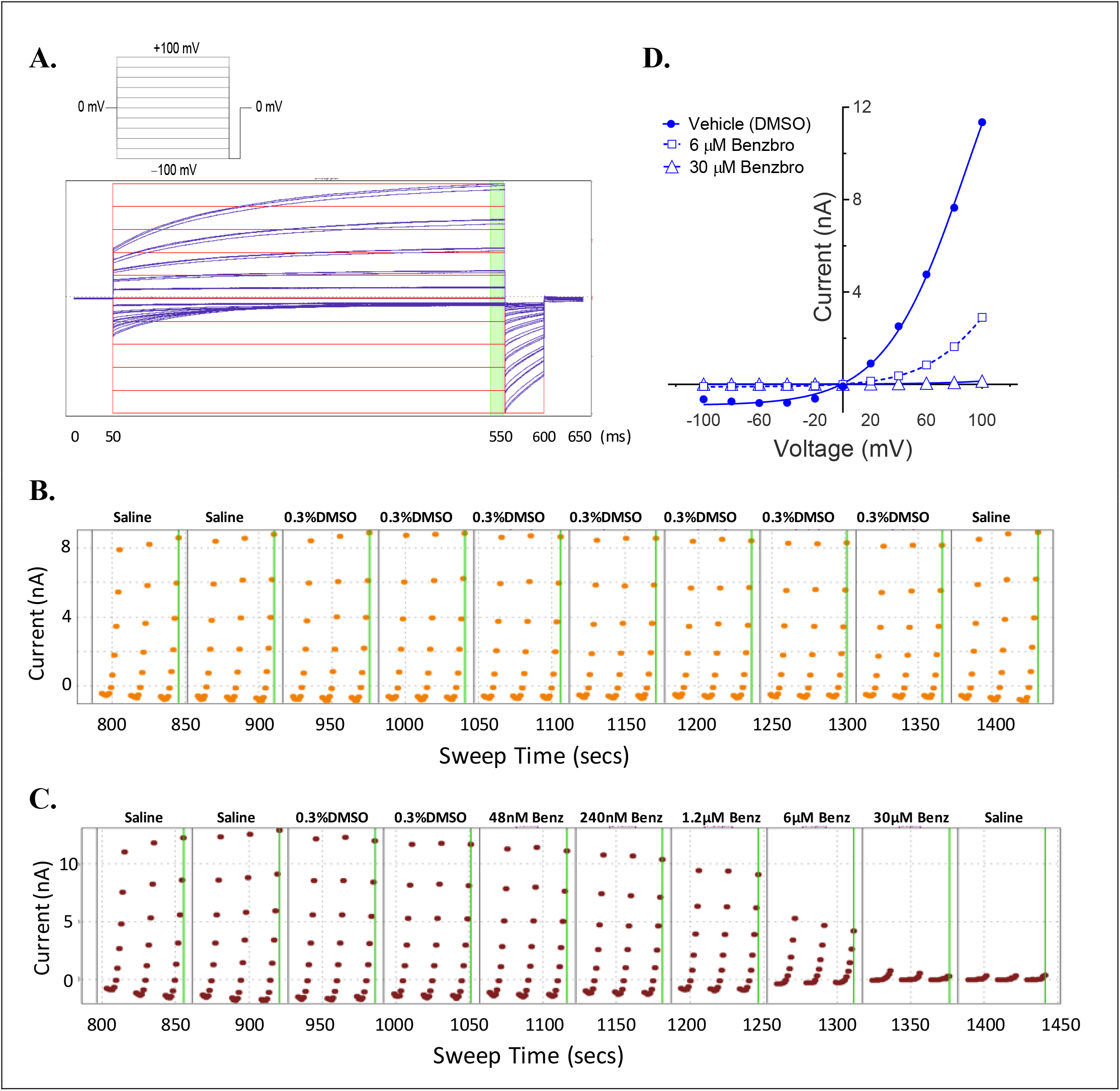
A medium-throughput, optimized planar patch clamp electrophysiology assay for stable recordings and routine measures of compound activity in blocking TMEM16A. (**A**) Representative whole-cell currents of a HEK293 cell stably-transfected with TMEM16A (acd) using an intracellular solution containing 170 nM free calcium. The standard QPatch voltage protocol had cells clamped from a holding potential of 0 mV (50 ms) to voltages between −100 and +100 mV in 20 mV steps (500 ms) followed by a step to −100 mV for 50 ms. There were 1.2 s between sweeps, 20 s between protocols, and 3 protocols run per sample addition which allowed at least 60 s incubation time following each vehicle or compound addition. (**B**) The TMEM16A currents were stable for at least 10 min and after repeat additions of the vehicle control (0.3% DMSO) in the optimized assay as shown in the representative recording, where the dots reflect the measured current from 540-550 ms after each 20 mV step from −100 to +100 mV, and three IV protocols are shown for each sample addition. Benzbromarone caused a dose-dependent inhibition of TMEM16A, which exhibited the typical outward rectification as shown in the representative instrument recording in (**C**) and the IV curves plotted in (**D**) of data from a separate recording showing currents measured at each voltage step during the third IV protocol.

Niclosamide, nitazoxanide and related compounds were found by QPatch electrophysiology to be potent inhibitors of the TMEM16A Ca^2+^-activated Cl^−^ current (**Figure 4A**). Significantly, while these compounds and 1PBC only partially inhibited the iodide/eYFP response (**Figures 1H,I**), they provided nearly complete inhibition of the *chloride* current as shown in the bar plots in **Figure 4B** and the dose-response curves of **Figures 4C** and **4D**. Over 80% of the calcium-activated chloride current was inhibited by niclosamide, Compounds 2-5, nitazoxanide and 1PBC, which was a similar scope of block to that observed with the benchmark antagonist, CaCCinh-A01. This stands in stark contrast to the results from the TMEM16A/eYFP assay using iodide as the permeant anion where the maximal percent inhibition by niclosamide of 62.4 ± 12.0 (n=39), as well as related compounds, was considerably less than that of CaCCinh-A01 which provided 102.1 ± 9.5 (n=53) percent inhibition (**Figure 1I**). This would suggest their efficacy depends on the permeate anion, with there being much greater impact in blocking chloride versus iodide permeation. The pharmacology of niclosamide and related compounds, like 1PBC, also contrasts with the benchmark antagonist benzbromarone, which fully inhibited both the iodide eYFP response and the TMEM16A chloride current. Importantly, however, 1PBC, niclosamide and related compounds, Cpd 2 and Cpd4, fully inhibited the TMEM16A calcium-activated chloride current yet were inactive in blocking the cAMP-induced CFTR chloride current as measured by IonWorks Barracuda (IWB) electrophysiology (**Figure 4E**; Supplementary Figure 26). In contrast, the benchmark inhibitors benzbromarone, CaCCinh-A01 and MONNA were less selective as they reduced both TMEM16A and CFTR chloride currents (**Figure 4E**; Supplementary Figure 27).

**Figure 4.**
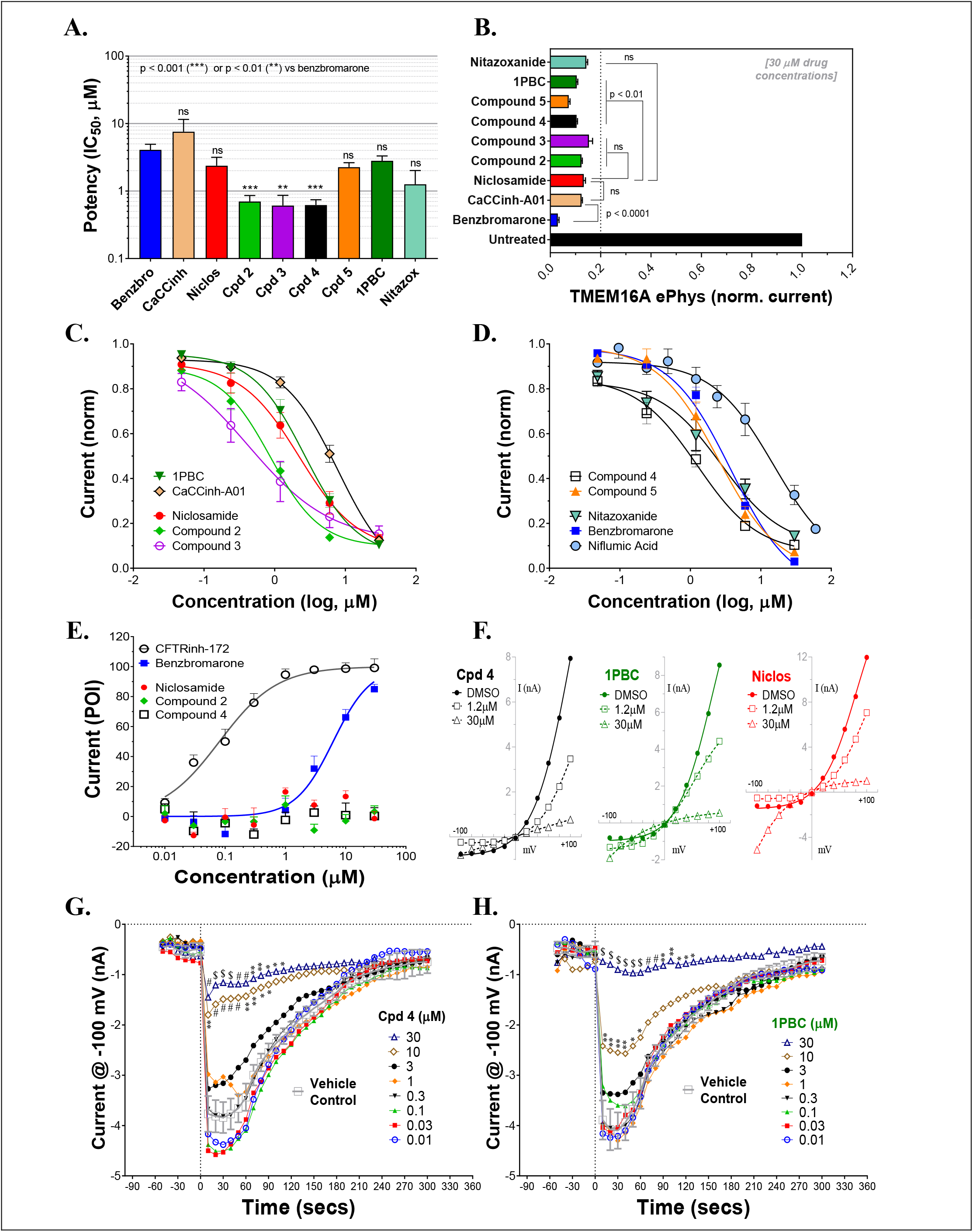
Niclosamide and related compounds provide potent and nearly full inhibition of TMEM16A chloride currents. Whole cell patch clamp studies using 170 nM free intracellular calcium revealed the average potency of TMEM16A antagonists in blocking the TMEM16A outward current at +100 mV (**A**), the compounds maximum antagonist activity (**B**) and concentration-response relationship for inhibition (**C, D**). Niclosamide, Cpd 2 and Cpd4 showed potent inhibition of the TMEM16A calcium-activated chloride current (**A - D**) but spared the cAMP-induced CFTR chloride current (**E**) exhibiting < 20 percent of inhibition (POI) of the forskolin-induced current. Panel (**F**) shows representative current-voltage traces from TMEM16A whole cell patch clamp electrophysiology studies using 170 nM free intracellular calcium, where the y-axis is current in nA and the x-axis is membrane potential changes from −100 mV to +100 mV in 20 mV steps from Vhold of 0mV. While benzbromarone inhibited both the outward and inward currents (Fig. 3), Cpd 4, 1PBC and niclosamide suppressed the outward current but showed concentration-dependent effects on the inward current ranging from inhibition to stimulation (**F**). The stimulation was especially noticeable for niclosamide and 1PBC at −100 mV and higher concentrations. In contrast, 1PBC (**H**) and Cpd 4 (**G**) were found to provide dose dependent inhibition (not stimulation) at −100 mV when tested by perforated patch clamp electrophysiology, where the ionomycin stim (10 μM) and compounds (from 0.01 – 30 μM) were added at the 0 sec time point. The DMSO Vehicle Control (grey, open squares; mean±SD, n=8) included on each patch plate shows the rapid calcium-dependent activation and inactivation of TMEM16A described earlier. ** P < 0.01, *** P < 0.001; significantly different from benzbromarone (**A, B**) or niclosamide (B); ns, not significant (unpaired *t*-test). Panels (**G**) and (**H**), P<0.05 (*), P<0.01 (**), P<0.001 (#), p<0.0001 ($) from two-way ANOVA for differences from the Vehicle Control; all other data points showed no significant difference (p>.05) compared to Vehicle. Number of replicates: n=3-11(**A-D**), n=3-4 (**E**), mean±SEM. Average of quadruplicate measures (**G, H**).

**Figure 5.**
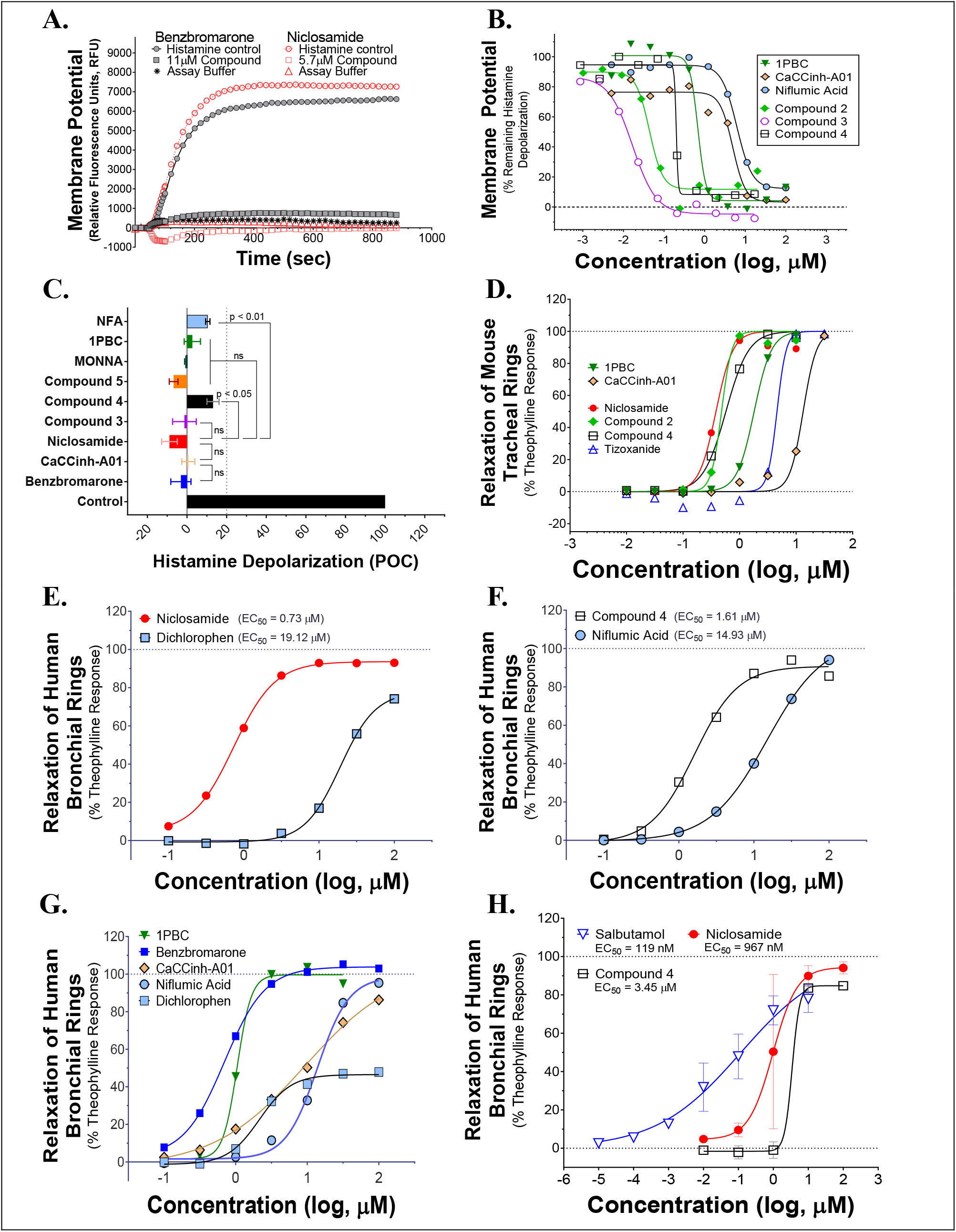
Niclosamide and related compounds block histamine depolarization of airway smooth muscle cells and provided robust bronchodilation of carbachol or histamine pre-contracted mouse and human airways. Representative kinetic traces of fluorescence recordings from the FLIPR membrane potential assay using primary human ASM cells (**A**). Histamine-induced a robust increase in fluorescence corresponding to depolarization which was fully blocked by the TMEM16A antagonists benzbromarone and niclosamide (**A**). Niclosamide and related compounds provided potent inhibition of the histamine-induced ASM depolarization (**B**), with the average maximum inhibition from dose-response studies exceeding 80% or <20 percent of control (POC) response to vehicle (Control); n = 2-15 (**C**). In wire myograph assays using carbachol pre-contracted mouse tracheal rings, niclosamide and related analogs, Cpd 2 and Cpd 4, provided sub-μM bronchodilation and improved potency over the two TMEM16A benchmark antagonists run in parallel (**D**). Niclosamide and Compound 4 also provided robust relaxation of human bronchial rings pre-contracted with carbachol, with results in panels (**E**) and (**F**) being wire myograph studies using rings from the same donor and involving simultaneous tests in parallel of the four compounds shown. Using a separate donor, effective bronchodilation of carbachol pre-contracted human airway rings was also observed with five benchmark TMEM16A antagonists of distinct chemotypes (**G**). Niclosamide and Compound 4, additionally provided robust relaxation of histamine pre-contracted human bronchial rings (n of 2-3) as shown in the wire myograph results in panel (**H**) on airway rings from different donors. The results from separate myograph studies on the β-agonist salbutamol are overlaid for comparison (n of 3 donors). As a precaution, all wire myograph studies using human bronchial rings included 5 μM indomethacin in the bath solution to eliminate any possible indirect effects due to prostaglandins. Mean±SEM. Significant differences from benzbromarone or niclosamide (**C**) was determined with unpaired *t*-test (ns, not significant).

**Figure 6.**
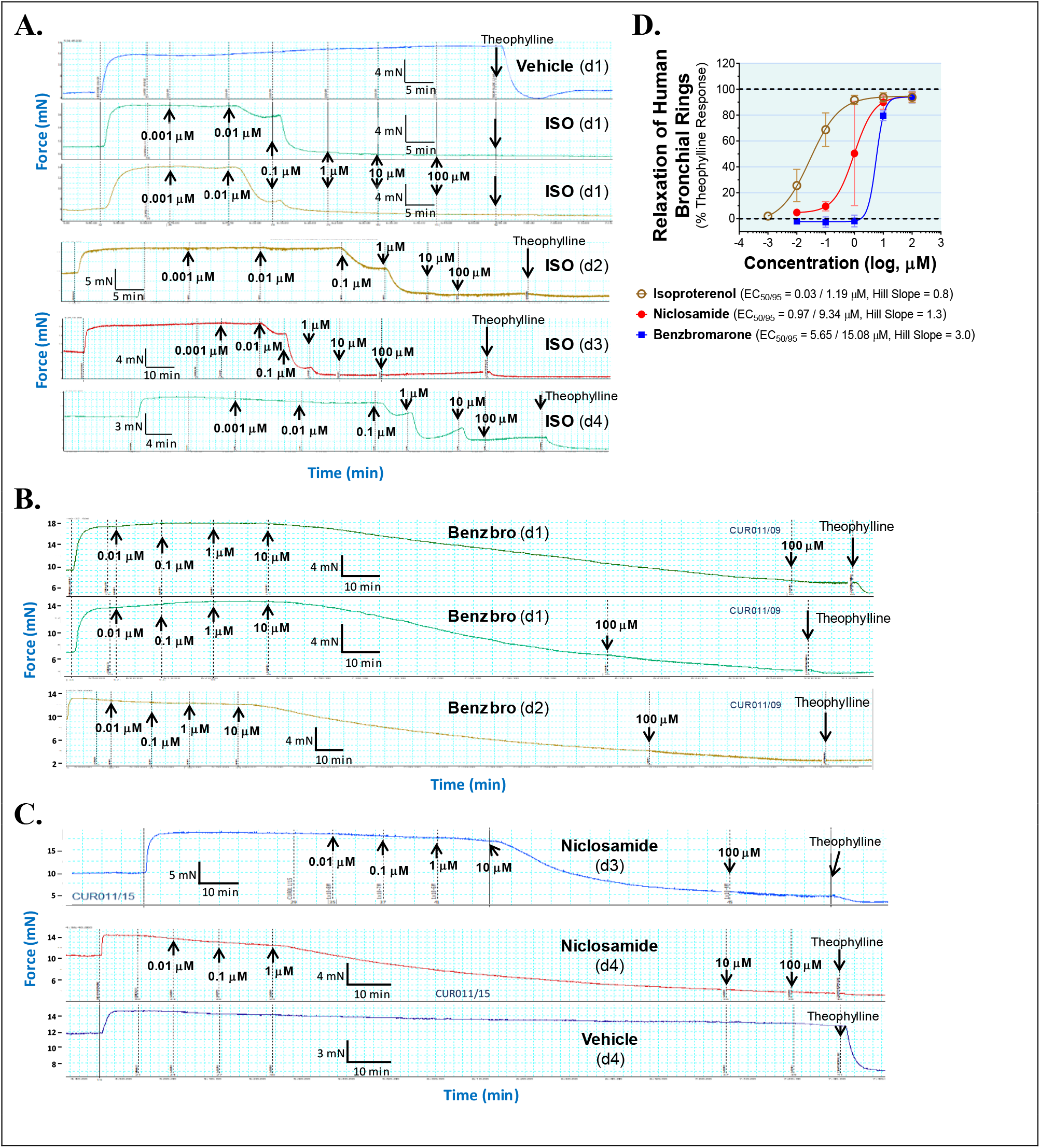
The hydrophilic β-agonist, isoproterenol, acts quickly to relax human bronchial rings while the lipophilic TMEM16A antagonists, benzbromarone and niclosamide, have a slow onset of action. Raw wire myograph recordings of force changes over time are shown for isoproterenol **(A)**, benzbromarone **(B)**, and niclosamide **(C)** relaxation of human bronchial rings pre-contracted with histamine and the average concentration-response relationship **(D)**. Force changes after histamine addition are stable over time and unaffected by repeat additions of Vehicle control **(A** – top panel and **C** – bottom panel; vertical black dotted lines). Isoproterenol (ISO) is hydrophilic (XLogP3 = −3.49) and has a quick onset of action, taking less than ~ 7 min at 10 nM or 100 nM concentrations to achieve steady-state relaxation of the human bronchial rings **(A)**, with results from four donors (d1-d4) being shown. In contrast, the lipophilic TMEM16A antagonist benzbromarone (Benzbro) (XLogP3 = 5.66) has a slow onset of action **(B)** in relaxing bronchial rings from donors d1 and d2, taking 80-140 min at 10 μM before force changes have reached a plateau. Niclosamide which is also lipophilic, XLogP3 = 5.32, similarly acted slowly taking ~ 50 min at 10 μM and ~ 92 min at 1 μM to achieve steady-state relaxation of the bronchial tissue from donors d3 and d4 **(C)**. To confirm responsiveness of each donor, isoproterenol is routinely run as a positive control. Mean ± SEM, N = 2-5.

Compounds 2–4 were more potent inhibitors of the calcium-activated chloride current than the benchmark antagonist benzbromarone (**Figure 4A**). This finding is consistent with results from the TMEM16A eYFP assay (**Figure 1G**). Niclosamide, however, which stood out as the most potent compound in the halide-sensitive YFP assay with average IC_50_ of 0.132 ± 0.067 μM (n=42), showed significantly less activity by QPatch electrophysiology with an average IC_50_ of 2.37 ± 2.11 μM (n=7) (**Figure 4A**). A consideration of the two TMEM16A assays may provide an explanation for these differences. In the high-throughput eYFP assay there is a 30 minute pre-incubation with compounds prior to ionomycin activation of the channel, while the automated QPatch electrophysiology assay has only about 1 min incubation time following each compound addition in the cumulative concentration response analyses (**Figure 3**). If there is insufficient time for the compound to block, one might expect to see currents still decreasing between the first and third I/V traces after each compound addition indicating steady-state block was not achieved. Supplementary Figures 6-21 provide representative QPatch instrument current-time recordings for niclosamide, nitazoxanide and related compounds, as well as, seven benchmark antagonists. While benzbromarone acts quickly (**Figure 3C** and Supplementary Figure 6) to block TMEM16A currents, we routinely observe currents continue to decline between the 1st and 3rd IV protocol after niclosamide addition (Supplementary Figure 12) indicating steady-state block is not achieved and its potency may be underestimated. Indeed, if we modify the protocol to look at single concentrations of niclosamide incubated for the maximum 12-15 minutes where we see stable recordings, we find for 1.11 μM niclosamide it took about 10 min to achieve steady state block (Supplementary Figure 13A) while approximately 4 min are needed when using the higher concentration of 3.3 μM (Supplementary Figure 13B). Similar studies on Compounds 2-4 (Supplementary Figure 14-18) indicate they also act slowly, and their potency may be underestimated by our automated QPatch assay. Unfortunately, extending the incubation time after each compound or vehicle addition resulted in too much current run-down and use of just single concentrations per recording chamber was too limiting to assay throughput. Despite these technical limitations, the electrophysiology data in **Figures 4A-4D** provided important evidence that niclosamide and related compounds were potent inhibitors of the TMEM16A Ca^2+^-activated Cl^−^ current.

As mentioned earlier, electrophysiology studies on TMEM16A are hampered by a strong calcium-dependent inactivation so we employed low intracellular calcium (170 nM free) and measured compound impact in blocking the outward current at +100 mV as little inward current is generated (**Figure 3**). Under physiological conditions, however, pro-contractile agonists would significantly increase intracellular calcium to the micromolar range resulting in a robust and biologically relevant TMEM16A inward current leading to depolarization and contraction of ASM cells. Interestingly, while only minor inward currents are observed when using low intracellular calcium and a holding potential of 0 mV, we found 1PBC and niclosamide caused a paradoxical increase in the inward current at −100 mV while robustly inhibiting the outward current in whole cell patch clamp electrophysiology studies (**Figure 4F**, Supplementary Figures 11-13). The benchmark TMEM16A antagonist, niflumic acid, was also reported earlier to show such voltage-dependent effects on TMEM16A (Bradley et al. 2014, Liu et al. 2015). We have confirmed these results, showing that niflumic acid, just like niclosamide, is blocking the outward yet stimulating the inward current (Supplementary Figure 10). The benchmark antagonist NTTP showed a similar pharmacology (Supplementary Figure 19), while the niclosamide-related compounds, Cpd 3 and Cpd 4 inhibited the outward current but showed no significant effect in either stimulating or inhibiting the inward current (**Figure 4F**, Supplementary Figures 16 and 18). In contrast, the benchmark antagonists benzbromarone, CaCCinh-A01, and MONNA inhibited both the outward and inward current (**Figure 3D**, Supplementary Figures 6-8).

While our recording conditions were similar to earlier whole cell patch clamp studies on TMEM16A, we were concerned that the non-physiological conditions of symmetrical chloride, low intracellular calcium and large hyperpolarization steps from 0 mV to −100 mV could be contributing to these paradoxical effects. Perforated patch clamp electrophysiology offers an advantage over whole cell patch clamp as it preserves many of the intracellular components of the cell. We therefore examined a subset of compounds for effects in blocking the inward current at −100 mV following ionomycin-activation of the TMEM16A chloride current. IonWorks Barracuda perforated patch clamp electrophysiology studies revealed a large TMEM16A inward current following ionomycin-activation of the channel, which rapidly inactivated (**Figures 4G,H**). 1PBC (30 μM) showed robust inhibition of this inward current (**Figure 4H**) which was in stark contrast to the effects observed by whole cell patch clamp studies (**Figure 4F**) where instead a slight stimulation was observed. Similarly, niflumic acid, which caused a dose-dependent stimulation of the inward current by whole cell patch clamp analysis (Supplementary Figure 10) instead by perforated patch clamp electrophysiology showed inhibition (not stimulation) of the inward current (Supplementary Figure 25C). These findings are consistent with the physiological effect of niflumic acid in blocking agonist-evoked and spontaneous-transient inward currents in airway smooth muscle (Liu and Farley 1996, Gallos et al. 2013). In contrast, the benchmark TMEM16A antagonist benzbromarone was effective in blocking the inward currents in both assays (**Figure 3D**, Supplementary Figure 25B).

It seems clear from these studies that 1PBC, niclosamide and related compounds (including several hundred not described here; see Supplementary Figure 4) exhibit a distinct pharmacology from the benchmark antagonist benzbromarone. To further assess the physiological effects of TMEM16A antagonists, we evaluated their efficacy in modulating airway smooth muscle depolarization and contraction following treatment with pro-contractiles.

### TMEM16A antagonists block ASM depolarization and bronchodilate airways

Pro-contractiles such as histamine and methacholine signal through Gq-coupled GPCRs on the cell surface to cause membrane depolarization and contraction of airway smooth muscle cells. To enable high-throughput measures of the impact of TMEM16A antagonists on ASM responses, we developed a membrane potential assay using primary human ASM cells. As demonstrated in the FLIPR-Tetra kinetic traces shown in **Figure 5A**, histamine causes a rapid increase in fluorescence in the membrane potential assay related to ASM depolarization that is fully suppressed by niclosamide and the TMEM16A antagonist, benzbromarone. **Figure 5B** provides representative dose-response results on six TMEM16A antagonists indicating they fully inhibited the depolarization induced by an EC_90_ amount of histamine (Supplementary Figure 28) and provided similar maximum inhibition compared to benchmark antagonists (**Figure 5C**). The niclosamide-related compounds, Cpd2 - Cpd4, exhibited significantly greater potency than the benchmark antagonists CaCCinh-A01 and niflumic acid in blocking ASM depolarization (**Figure 5B**). Heartened by these results we advanced compounds to tissue studies using mouse tracheal rings to evaluate their effect in bronchodilating airways. As shown in **Figure 5D**, niclosamide, Cpd2 and Cpd 4 were highly potent (sub-μM) in providing full relaxation (bronchodilation) of carbachol precontracted airways. Tizoxanide, the metabolic product of nitazoxanide, also provided efficient bronchodilation of mouse airways. **Table 1** provides the full results from studies on these compounds and additional TMEM16A antagonists, where it can be seen the enhanced efficacy of niclosamide and related compounds in blocking TMEM16A corresponds to improved potency in bronchodilating airways. For the first time, we also provide data exploring the effects of 1PBC on ASM physiology. While 1PBC to date has only been explored for effects on TMEM16A, our data would suggest it is highly effective in blocking depolarization and contraction of ASM cells (**Figures 5B,D**). NTTP, which is structurally related to tizoxanide (**Figure 2**), was also efficacious in relaxing mouse tracheal rings (Supplementary Figure 36). For over 50 years there has been no new bronchodilators, with β-agonists remaining the only agent and mechanism for blocking the multiple contractiles operating in disease. Antagonists of TMEM16A offer a new mechanism, thus it’s important to point out our studies with ASM cells using histamine and our studies with mouse trachea using the cholinergic carbachol support the idea that TMEM16A antagonists block the multiple contractiles that can operate in disease. To extend our bronchodilation studies to human tissue and smaller airways, we’ve also tested antagonists for relaxation of human 4th order bronchi. Niclosamide and Compound 4 fully relaxed human bronchial rings pre-contracted with carbachol (**Figures 5E,F**) or histamine (**Figure 5H**). Of the TMEM16A antagonists, niclosamide routinely provided the most potent bronchodilation. In general, its efficacy in relaxing human bronchi compared favorably to its efficacy in relaxing mouse tracheal rings where it averaged an EC_50_ of 0.379 μM (**Table 1**). Of the benchmark TMEM16A antagonists, 1PBC had the greatest potency in relaxing carbachol precontracted human bronchial rings with EC_50_ of 0.91 ± 0.20 μM, followed by benzbromarone, CaCCinh-A01, dichlorophen and niflumic acid with EC_50_ values of 2.08 ± 1.95, 9.24, 10.64 ± 12.0 and 14.40 ± 0.76 μM, respectively (**Figures 5E-G**, Supplementary Figure 38). The short-acting β-agonist salbutamol had an EC_50_ of 119 nM for bronchodilating histamine precontracted human airways that was significantly lower than the EC_50_ of niclosamide (967 nM) and Cpd 4 (3.45 μM), but a similar concentration of ~ 10 μM was needed to fully relax the tissue (**Figure 5H**).

### TMEM16A antagonists have a slow onset of action, but could offer sustained effects in bronchodilating airways

The onset and duration of bronchodilation are two additional variables to explore when developing new asthma therapies. These can be examined with wire myograph by assessing force changes over time. The raw wire myograph traces of force changes over time for isoproterenol, benzbromarone, and niclosamide relaxation of human bronchial rings and the average concentration-response relationships are shown in **Figure 6**. The hydrophilic β-agonist isoproterenol (XLogP3 = −3.49) had a quick onset of action, relaxing the histamine pre-contracted human bronchial tissue in less than 7 min at 10 or 100 nM doses (**Figure 6A**). In contrast, the lipophilic TMEM16A antagonists benzbromarone and niclosamide (XLogP3 of 5.66 and 5.32, respectively) had a slow onset of action in relaxing human bronchial rings pre-contracted with histamine. Benzbromarone took 80-140 min at a 10 μM dose (**Figure 6B**), while niclosamide required ~50 min at 10 μM and ~92 min at a 1 μM before force changes reduced to a plateau (**Figure 6C**). Compound 4 is also lipophilic (XLogP3 = 4.63) and took between 50 – 125 min at 10 μM in three separate wire myograph recordings before fully relaxing the tissue (Supplementary Figure 40). Despite the significant difference in onset of action between the β-agonist isoproterenol and the TMEM16A antagonists, all compounds fully relaxed human bronchial rings pre-contracted with histamine (**Figure 6D**).

Similar to the effects on human bronchial rings, benzbromarone, niclosamide and Cpd 4 had a slow onset of action in relaxing mouse tracheal rings pre-contracted with carbachol (**Figure 7**). Benzbromarone had the slowest onset of action taking 40 – 60 min at 1 μM to achieve steady-state relaxation of the tissue (**Figure 7D**), while niclosamide took 17 – 35 min at the same dose (**Figure 7E**) and Cpd 4 required 25 – 45 min at 316 nM (**Figure 7F**) before relaxation had reached a plateau. As was the case with human bronchial rings, the hydrophilic β-agonist isoproterenol acted quickly to relax mouse airways taking just 2 – 4 min to relax the tissue (Supplementary Figure 39). For the lipophilic TMEM16A antagonists, a technical limitation in wire myograph experiments due to the slow onset of action is a tendency to overlook activity at lower concentrations where insufficient time is allowed to relax the tissue. We think this can lead to compression of the dose-response curves and noticeably large Hill Slopes, such as those observed in **Figures 7A-D**.

**Figure 7.**
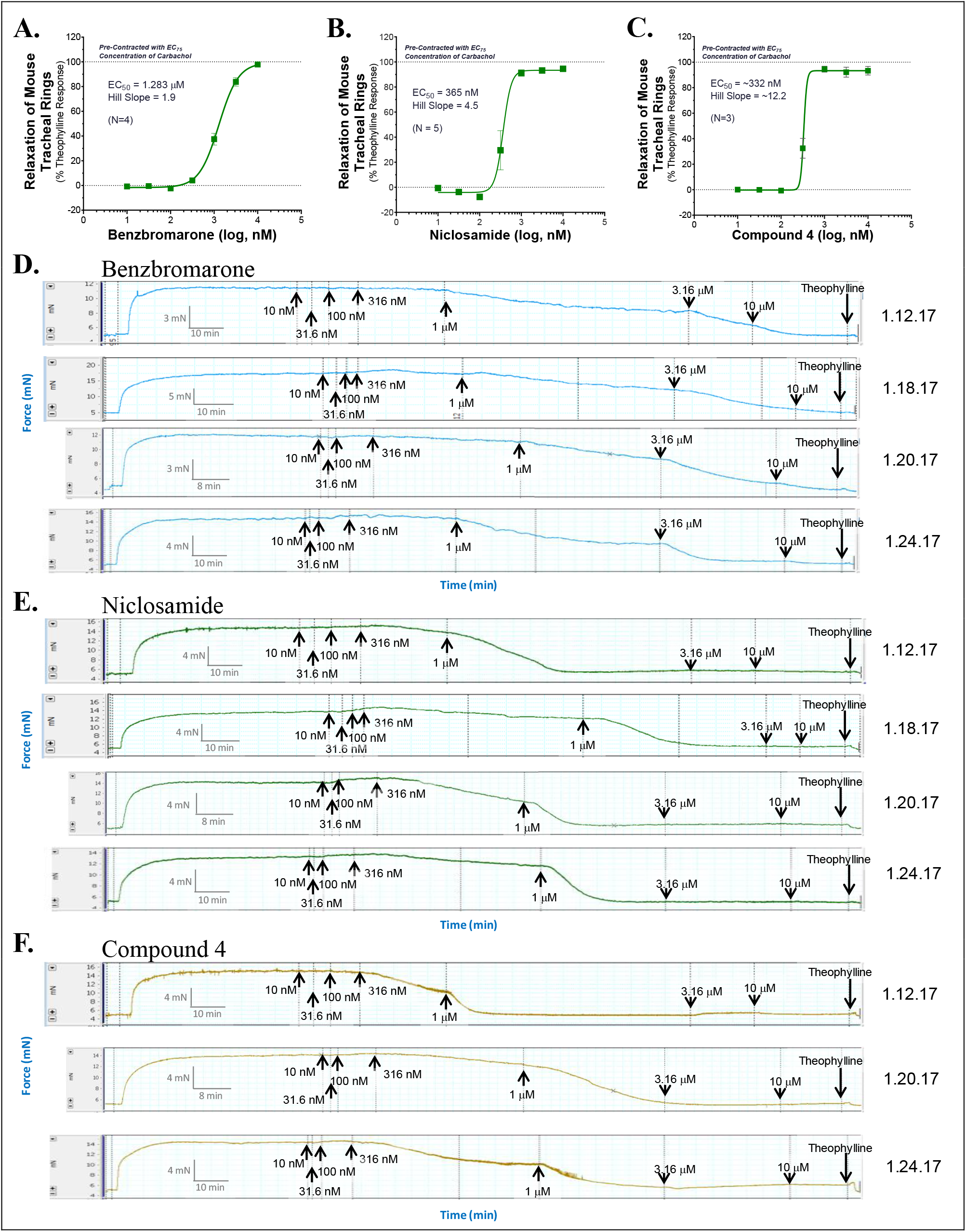
The TMEM16A antagonists, benzbromarone, niclosamide, and Compound 4 fully relax mouse tracheal rings but have a slow onset of action. The average concentration-response relationship (**A-C**) and raw wire myograph recordings of force changes over time are shown for benzbromarone (**A, D**), niclosamide (**B, E**), and Compound 4 (**C, F**) for relaxation of mouse tracheal rings pre-contracted with an EC_75_ concentration of carbachol. The lipophilic compounds benzbromarone (XLogP3 = 5.66), niclosamide (XLogP3 = 5.32), and Cpd 4 (XLogP3 = 4.63) took an extended period of time to achieve steady-state relaxation of the tissue but had efficacy similar to theophylline in fully relaxing the mouse airways. At a 1 μM dose, benzbromarone (**D**) exhibited a slow onset of action taking between 40 – 60 min to achieve steady-state relaxation of the tissue, while niclosamide took between 17 – 35 min at the same dose (**E**). Cpd 4 also had a slow onset of action taking between 25 – 45 min at 316 nM to achieve steady-state relaxation (**F**). In contrast the hydrophilic β-agonist isoproterenol (XLogP3 = −3.49), while only partially relaxing airway rings pre-contracted with an EC_75_ level of carbachol, acted quickly taking just 2 – 4 min to bronchodilate the tissue (Supplementary Figure 39). To enable direct comparison of the activity of the molecules, compounds were run in parallel on the same day in separate chambers of the wire myograph. Theophylline was routinely added at the end of the recording to define full bronchodilation and calculate percent relaxation of the tissue. The results shown are from 3-5 separate experiments. Mean ± SEM.

To explore compound duration of action, we treated tissue with compounds, washed and then re-challenged the tissue with pro-contractiles to see if bronchodilation was sustained or transient. Preliminary tests of the β-agonist isoproterenol indicate its effects were short-acting as expected, while bronchodilation with niclosamide and benzbromarone were long lasting (Supplementary Figure 37).

### Antagonists resist use- and inflammatory-desensitization mechanisms

TMEM16A antagonists offer a new mechanism to bronchodilate airways that we’d expect to resist use- and inflammatory-desensitization mechanisms that limit β-agonist action. To test this hypothesis, we employed mouse tracheal rings which can be readily attained and compared TMEM16A antagonists versus the β-agonist isoproterenol for bronchodilation after various challenges meant to reflect the situation of severe asthma with poorly controlled inflammation and airway hyperresponsiveness (AHR), which reflects increased sensitivity to contractants. While β-agonists fully relax partially precontracted airways, it’s been shown earlier they only partially relax maximally precontracted airways (Lemoine and Overlack 1992), which may be relevant to drug therapy and disease control. We have confirmed this desensitization phenomena using the β-agonist isoproterenol, but importantly find TMEM16A antagonists fully bronchodilate airways irrespective of the level of carbachol precontraction (**Figure 8**). To accomplish these studies, the EC_25_, EC_50_, EC_75_ and EC_95_ for carbachol bronchoconstriction is first determined for each airway ring (Supplementary Figure 33), the tissue is then washed to return tissue to baseline tension and then treated with various levels of contractant. As shown in **Figure 8A**, the β-agonist isoproterenol fully (100%) relaxed airway rings treated with an EC_25_ level of carbachol but caused only a partial 42% relaxation of airways treated with an EC_95_ level of carbachol. In contrast, the TMEM16A antagonists benzbromarone, niclosamide and Compound 4 fully relaxed maximally (EC95) contracted airways, provided roughly 100% bronchodilation irrespective of contractile force (**Figures 8B-D**). Thus, TMEM16A antagonists as an add-on therapy operating with a distinct MOA may help patients retain disease control when β-agonists are functioning sub-optimally. Indeed, an airway ring maximally contracted with an EC_95_ level of carbachol showing just 38% bronchodilation with 1 μM isoproterenol, achieved roughly 100% bronchodilation after subsequent treatment with the TMEM16A antagonist, Cpd 4 (**Figure 8A**).

**Figure 8.**
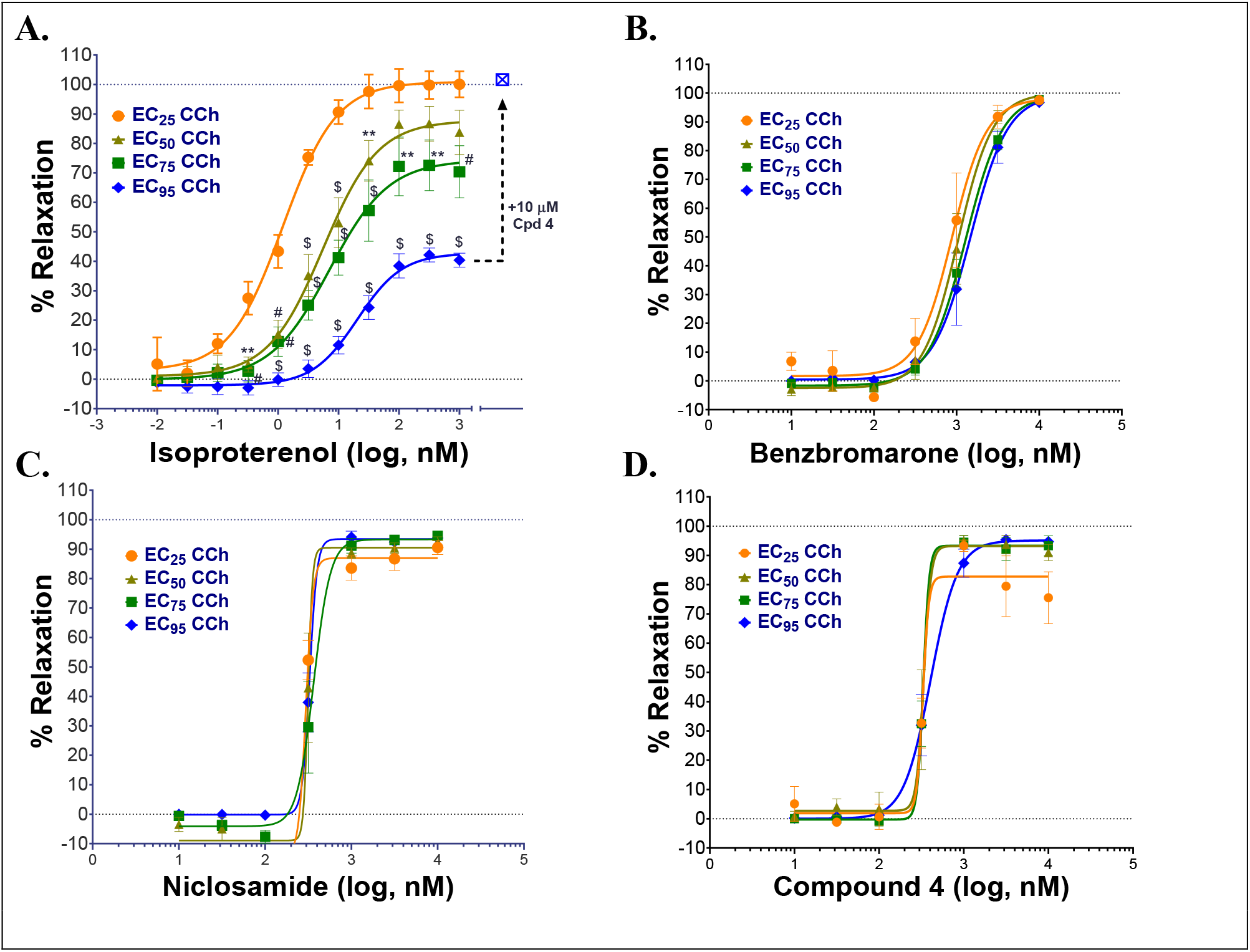
TMEM16A antagonists fully relax airways in the presence of low and high concentrations of contractants, while the β-agonist isoproterenol showed only limited efficacy on maximally contracted airways. Mouse tracheal rings were pre-contracted with EC_25_ – EC_95_ concentrations of carbachol (CCh) and then treated with the β-agonist isoproterenol **(A)**, or the TMEM16A antagonists benzbromarone **(B)**, niclosamide **(C)** or Compound 4 **(D)** to measure bronchodilation using a DMT wire myograph (n=3-5 rings per condition). A maximally contracted airway ring using an EC_95_ level of carbachol that showed only partial bronchodilation after the highest isoproterenol dose, was fully relaxed by add-on of the TMEM16A antagonist Cpd 4 (% relaxation shifts from 37.7% → 101.7% at EC_95_); see arrow to open blue square, cross-hatched **(A)**. Mean±SEM. P≤.05 (*), P≤.01 (**), P≤.001 (#), P≤.0001 ($), two-way ANOVA with Dunnett’s multiple comparison test for differences from the EC_25_ of CCh in each group. Only the isoproterenol group showed a significant difference in the dose-response relationship depending on the level of CCh pre-contraction, while no significant effects were observed for the benzbromarone, niclosamide or Cpd 4 groups.

Severe asthma and COPD is associated with poorly controlled inflammation. Using a cocktail of cytokines (IL-1β, TNFα, IL-13) to mimic this process, we show the efficacy of isoproterenol in bronchodilating airways is dramatically reduced when airways are pretreated with cytokines (**Figure 9A**). In contrast benzbromarone and niclosamide fully bronchodilated cytokine-treated airways (**Figures 9B,C**), suggesting TMEM16A antagonists offer a new mechanism to resist the inflammatory desensitization pathways that limit β-agonist action. Repeat treatment with β-agonists are also known to induce tachyphylaxis or use-dependent desensitization, which involves β2-adrenergic receptor phosphorylation, arrestin recruitment and receptor internalization and degradation resulting in reduced bronchodilation. To ascertain whether these pathways leading to β-agonist desensitization also interfere with bronchodilation by TMEM16A antagonists, we measured the degree of bronchodilation after repeated treatment with isoproterenol. As shown in **Figure 9D**, there was less bronchodilation and a rightward shift in potency after the 2nd round of β-agonist treatment which reflected use-dependent desensitization. Subsequent addition of Compound 4 to this same tissue resulted in full bronchodilation indicating TMEM16A antagonists restore bronchodilation of airways undergoing β-agonist use dependent desensitization (**Figure 9D**). The combined data indicates TMEM16A antagonists provide a new mechanism for robust relaxation of airway smooth muscle, even when they encounter high levels of contractants or are exposed to inflammatory conditions that limit β-agonist action.

**Figure 9.**
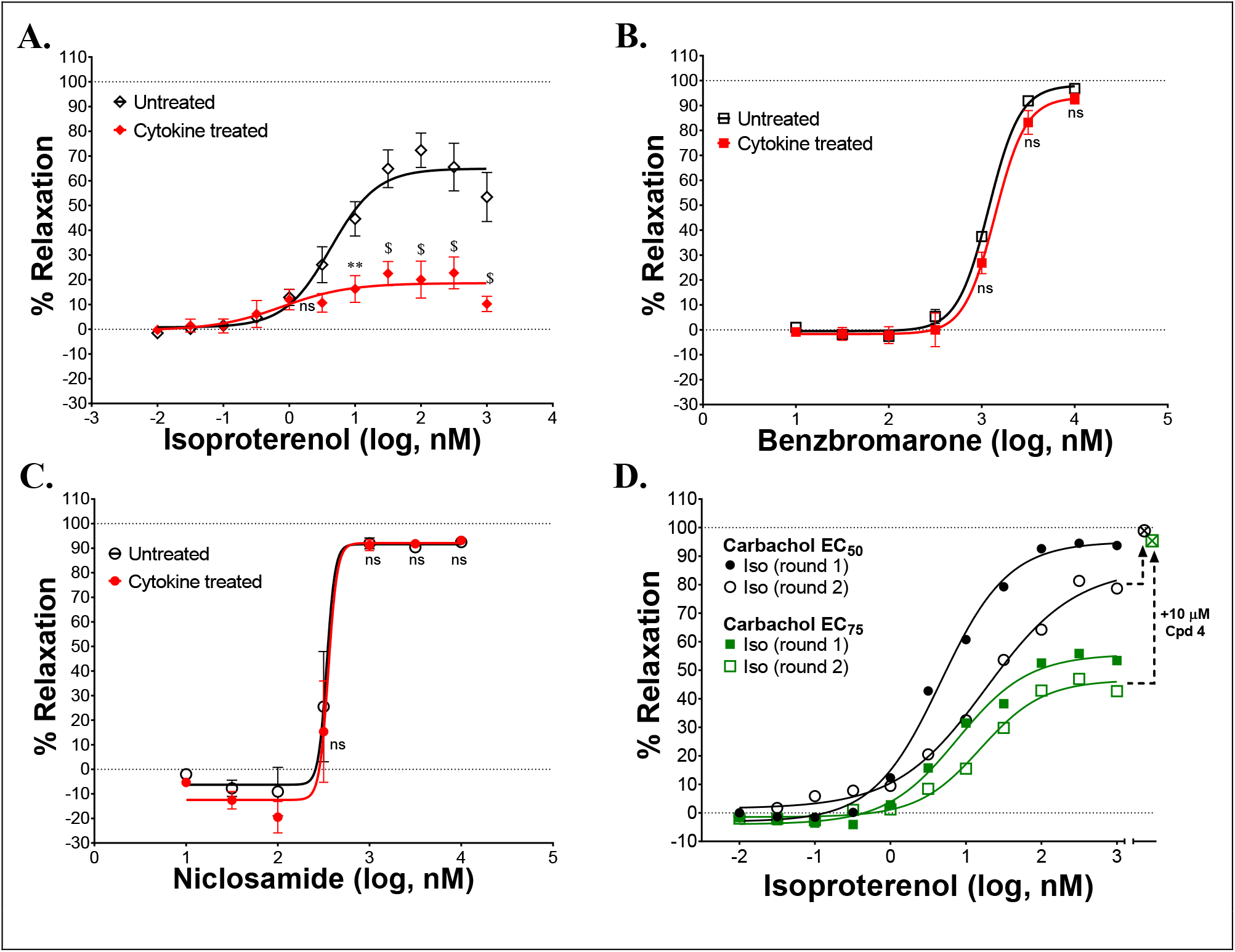
TMEM16A antagonists fully bronchodilate airways and resist use- and inflammatory-desensitization pathways that limit β-agonist efficacy. Treatment of mouse tracheal rings overnight with a cytokine cocktail of IL-1β, IL-13 and TNFα strongly reduced isoproterenol bronchodilation of carbachol pre-contracted airways **(A)**, while the TMEM16A antagonists benzbromarone **(B)** and niclosamide **(C)** remained effective in fully relaxing the cytokine treated airways (n=4; 2 rings/group, 2 experiments). As a model of β-adrenergic receptor use-dependent desensitization, carbachol pre-contracted (EC_50_ or EC_75_ level of CCh) mouse tracheal rings were treated with the β-agonist isoproterenol (round 1), washed, re-contracted with carbachol and tested again (round 2) for bronchodilation. Isoproterenol showed a rightward shift in potency after the second treatment with the β-agonist and a reduction in maximal efficacy in bronchodilation **(D)**. Airway rings showing desensitization and only partial bronchodilation after this second treatment and highest isoproterenol dose, were fully relaxed by add-on of the TMEM16A antagonist Compound 4 (% relaxation shifts from 78.6% → 98.9% at EC_50_ and from 42.7% → 95.4% at EC_75_); see arrows to open cross-hatched circle and square **(D)**. P≤0.05 (*), P≤0.01 (**), P≤0.001 (#), P≤0.0001 ($) from two-way ANOVA with Sidak’s multiple comparison test for differences between untreated and cytokine treated airway rings for each compound treatment group; all other data points showed no significant (ns) difference (p>0.05). Mean±SEM.

### TMEM16A expression in human ASM and bronchial epithelium, upregulation by Th2 cytokines and overexpression in lungs of monkeys with allergic asthma

Using an RNAseq dataset described earlier (Aisenberg et al. 2016), we evaluated TMEM16A expression in primary human airway smooth muscle cells and bronchial epithelial cells derived from healthy donors or patients with asthma or COPD. TMEM16A expression was higher in bronchial epithelial cells compared to ASM cells (**Figure 10A**), with the serial cultures of epithelial cells mostly comprised of basal cells. Expression was detected in both normal and diseased samples and confirmed at the sequence level by evaluating RNAseq read coverage (Supplementary Figure 41). Inflammatory cytokines can also act on the bronchial epithelium to induce goblet cell hyperplasia and mucin hypersecretion leading to airway narrowing and loss of lung function. To evaluate the effects of the Th2 cytokines, IL-4 and IL-13, on TMEM16A expression in the mature bronchial epithelium, we cultured normal and COPD human bronchial epithelial (HBE) cells at an air-liquid interface (ALI) to generate a polarized, pseudostratified bronchial epithelium composed of basal cells, secretory cells, and ciliated cells. The mature ALI cultures were then untreated or treated with IL-13 for 1, 3, 5 or 7 days and processed for RNAseq whole transcriptome sequencing to evaluate gene expression. TMEM16A was dramatically upregulated after IL-13 treatment of HBE ALI cultures quickly reaching a peak after just 1 day (**Figure 10B**, left panel), while elevated expression of the Muc5AC mucin occurred more slowly and progressively from days 3 to 7 (**Figure 10B**, right panel). Immunohistochemical analysis of separate ALI cultures, untreated or treated with IL-4 or IL-13 for 2 days confirmed TMEM16A expression is also increased at the protein level (**Figure 10C**). Increased staining was observed at the apical surface (air interface, right side of ALI) and at the basolateral surface near the transwell membrane insert (liquid interface, left side of ALI) of the mature bronchial epithelial ALI cultures after treatment with the Th2 cytokines (**Figure 10C**). In the latter case elevated TMEM16A staining appeared to be of basal cells, while in the former case it appeared there was increased staining of mucin secreting goblet cells following treatment with the Th2 cytokines. To evaluate TMEM16A expression in the lung of naive or asthmatic cynomolgus (cyno) monkeys, tissue was obtained from naïve adult cynos that were non-allergic to *Ascaris suum* allergen and from adult cynos exhibiting repeated sensitivity to *A. suum* allergen that were characterized as asthmatic. Significant increases in TMEM16A expression was detected at the apical surface of the bronchial epithelium and in submucosal glands of cynomolgus monkeys with allergic asthma compared to naïve non-allergic animals (**Figure 10D**). Immunohistochemistry of lung tissue from a separate monkey with allergic asthma localized the increased TMEM16A staining to the apical surface of goblet cells in the bronchial epithelium (**Figure 10E**).

**Figure 10.**
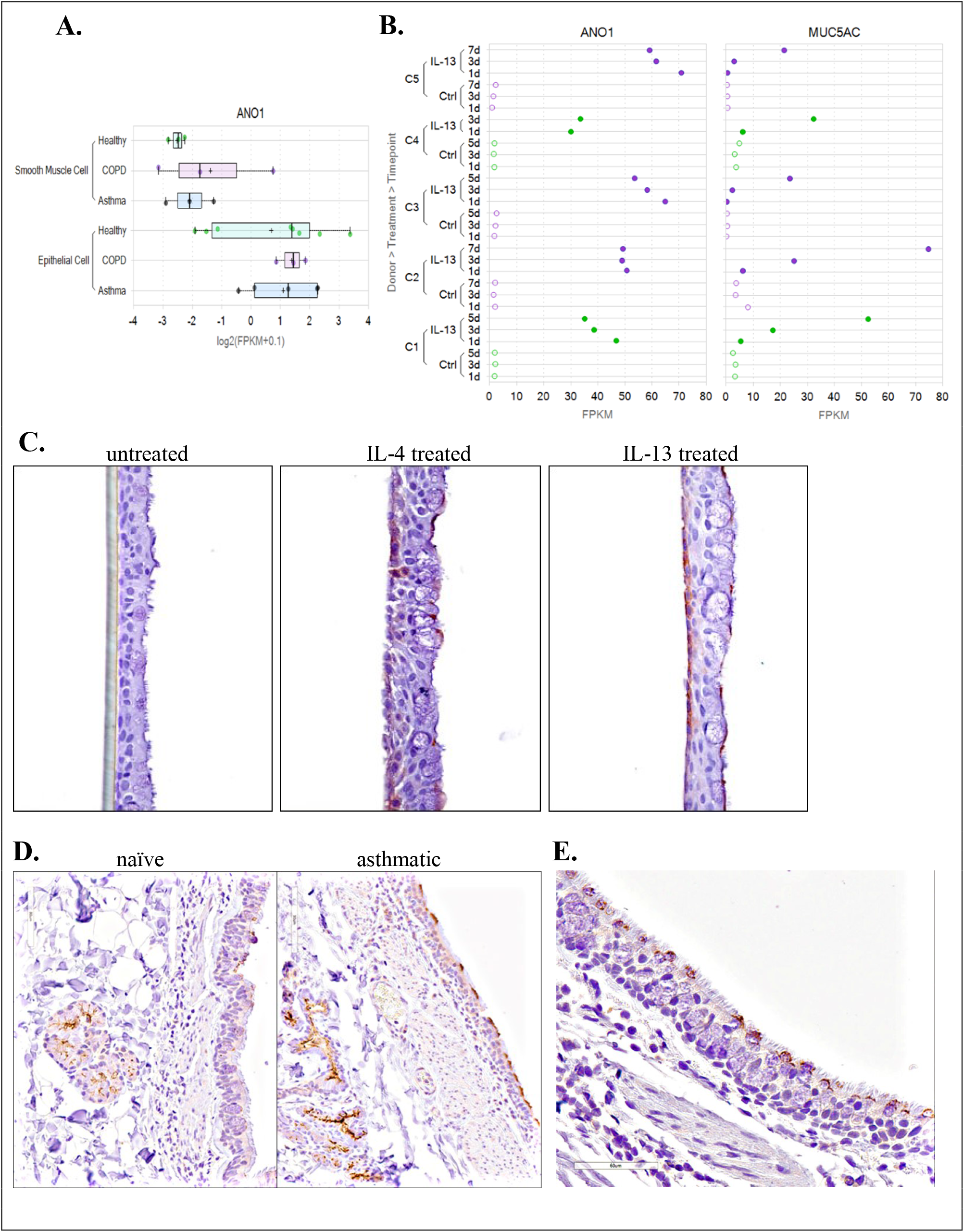
TMEM16A is expressed in cultured human airway smooth muscle and bronchial epithelial cells, quickly upregulated in mature human bronchial epithelial air-liquid interface (ALI) cultures after IL-4 or IL-13 treatment and overexpressed in the lung of asthmatic cynomolgus monkeys on the apical surface of goblet cells and in submucosal glands. TMEM16A (ANO1) expression in cultured human bronchial epithelial cells (HBE, submerged) and human airway smooth muscle cells from normal, asthmatic or COPD donors **(A)** was determined from an RNAseq dataset described earlier (Aisenberg et al. 2016). TMEM16A expression was higher in bronchial epithelial cells compared to airway smooth muscle cells. The results for ASM cells (normal or diseased) are from 3 separate donors each, while the results for HBE cells are the average response from 8 normal, 5 asthma and 3 COPD donors. The Th2 cytokine, IL-13, dramatically increased TMEM16A (ANO1) gene expression in mature bronchial epithelial ALI cultures reaching a maximum after 1 day **(B,** left panel), while elevated expression of the MUC5AC gene (B, right panel) occurred more slowly from day 1 through day 5 or 7. The RNAseq data shown are from ALI cultures from two normal (green; C1,C4) and three COPD (purple; C2, C3, C5) donors, untreated (Ctrl, open circles) or treated with IL-13 (closed circles). The Th2 cytokines, IL-4 and IL-13, also upregulated TMEM16A protein expression in human bronchial epithelial air-liquid interface cultures **(C)**. Fully differentiated human bronchial epithelial ALI cultures were untreated **(C,** left panel) or treated for 48 hours with IL-4 **(C,** middle panel) or IL-13 **(C,** right panel), fixed and stained with a rabbit anti-TMEM16A monoclonal antibody from Abcam (Ab64085). Increased TMEM16A staining (brown) was detected in basal cells located near the membrane insert and basolateral surface (left side of ALI culture) and on goblet cells located at the apical surface (right side) of IL-4 or IL-13 treated ALI cultures. Elevated TMEM16A protein expression was also detected in the bronchiolar epithelium and submucosal glands of cynomolgus monkeys with allergic asthma **(D)**. Cynomolgus monkeys repeatedly challenged with *Ascaris suum* were compared to naïve non-allergic monkeys for TMEM16A expression using the polyclonal antibody Ab53212 (Abcam). Increased TMEM16A staining (brown) was observed on the apical surface of the bronchial epithelium and in submucosal glands of monkey with allergic asthma **(D,** right panel; animal #384630) compared to a naïve non-allergic monkey **(D,** left panel; animal #384606). Lung specimens from a separate asthmatic cynomolgus monkey (#384750) were collected 24 hours post *A. suum* challenge. Immunohistochemical staining with the anti-TMEM16A antibody indicates TMEM16A is expressed selectively on the apical surface of goblet cells within the bronchial epithelium **(E)**.

### Antagonists effects on endogenous calcium-activated chloride currents and identification of TMEM16A as molecular target for niclosamide in cancer

TMEM16A is located within the 11q13 amplicon, one of the most frequently amplified chromosomal regions in human cancer that correlates with a poor prognosis. To validate the pharmacology of antagonists in modulating native calcium-activated chloride currents, we profiled RNAseq data on large panel of cancer cell lines to identify high expressers. Of the 110 cell lines showing the most elevated TMEM16A expression (Supplementary Figure 43), roughly 18 % were colorectal cancer cell lines and 6% were gastric cancer cell lines. We chose one of the colorectal cancer (CRC) cell lines, COLO205, for further evaluation by QPatch electrophysiology. COLO205 cells gave good seals and a robust TMEM16A current with the characteristic calcium and voltage-dependence. Application of three benchmark TMEM16A antagonists (benzbromarone, CaCCinh-A01 and 1PBC) fully blocked this native chloride current (**Figure 11A-C**; Supplementary Figures 22 and 23). Importantly, the paradoxical pharmacology observed earlier in electrophysiology studies on 1PBC using cloned TMEM16A in HEK293 (**Figure 4F**), was also observed in studies with COLO205 cells expressing native TMEM16A currents. To our knowledge this represents the first description that COLO205 cells express a functional TMEM16A channel, but interestingly the colorectal cell line, HT-29, was exploited in one of the first screens to identify antagonists of the calcium-activated chloride channel (De La Fuente et al. 2008) before its molecular identity was defined. While the confirmation of antagonist activity in blocking endogenous TMEM16A may seem trivial, notable differences have been observed between TMEM16A overexpressed in HEK293 cells and cell lines expressing TMEM16A endogenously (Schreiber et al. 2018), making it important to validate responses in both settings.

**Figure 11.**
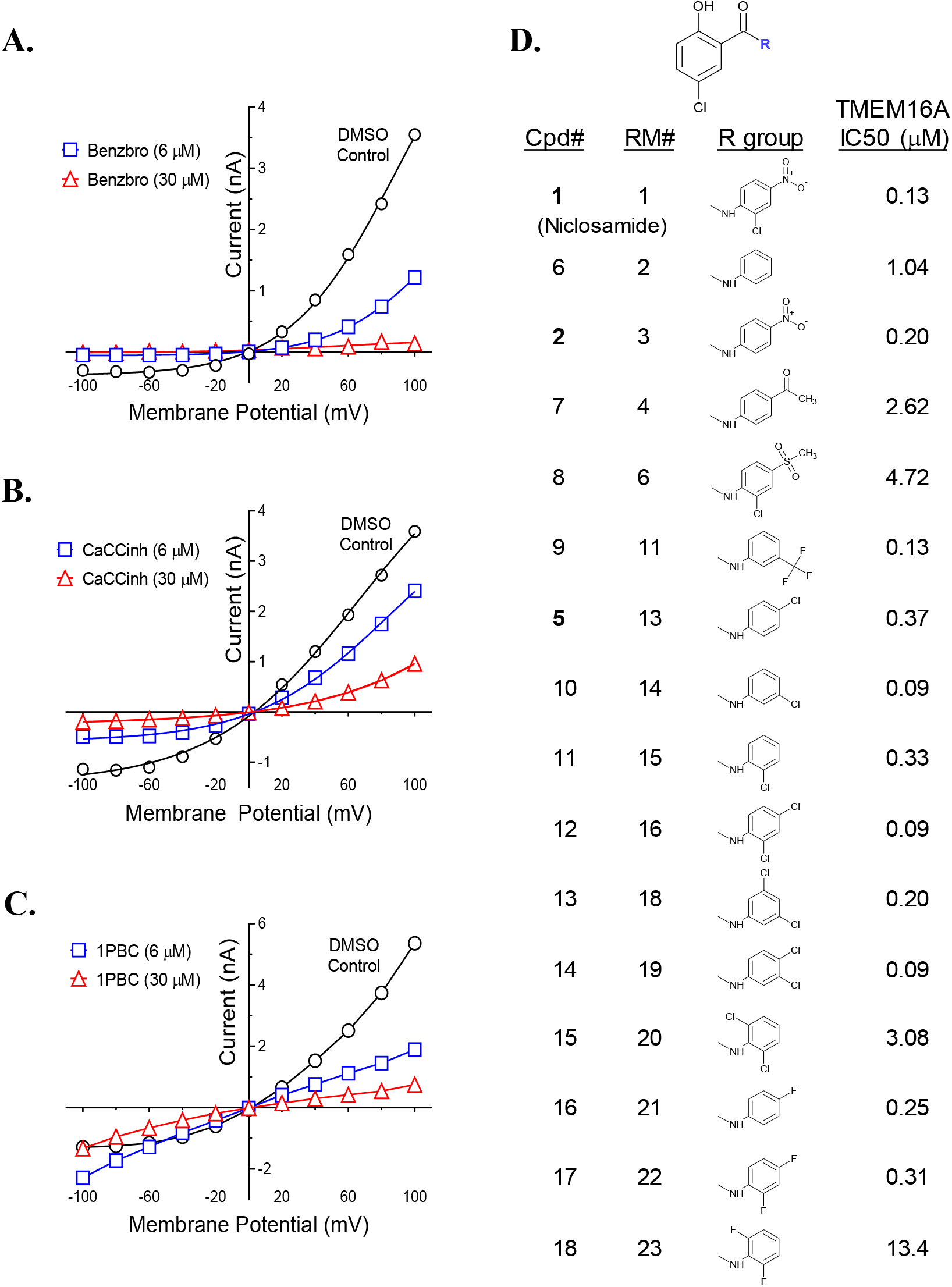
TMEM16A activity of additional niclosamide analogs and the impact of benchmark antagonists on endogenous calcium-activated chloride currents in the colorectal cancer cell line, COLO205. RNAseq identified several colorectal cancer cell lines as overexpressing TMEM16A (Supplementary Figure 43). Shown are results from QPatch electrophysiology studies indicating COLO205 cells exhibit the characteristic current-voltage relationship of TMEM16A which was sensitive to the TMEM16A antagonists benzbromarone **(A)**, CaCCinh-A01 **(B)** and 1PBC **(C)**, the later showing the same anomalous effects on the inward current observed with cloned TMEM16A in HEK293 cells. The structure activity relationship of several other niclosamide analogs is provided in **(D)**. Drug repurposing has identified niclosamide as a promising new treatment for cancer, but its mechanism is not fully understood. Mook et al. (2015) have found niclosamide and related molecules suppress signaling and proliferation of cancer cells, including from the colon cancer cell line HCT-116. We find these compounds are also potent TMEM16A antagonists. Our compound numbering (Cpd#), the compound numbering by R. Mook et al. (RM#), and the calculated potency (IC_50_, μM) of the molecules in our TMEM16A halide-sensitive eYFP assay are shown **(D)**.

Drug repurposing is an attractive approach to reduce the time and cost of drug development as it exploits preexisting clinical data on the pharmacokinetics, pharmacodynamics and safety of approved compounds. From such efforts niclosamide has emerged as a promising anticancer agent (Li et al. 2014). It inhibits proliferation of the colon cancer cell lines HCT116, COLO205 and HT29 with an IC_50_ of 0.37, 0.98 and 4.17 μM, respectively, which falls within the range of potency we find for blocking TMEM16A conductivity (**Table 1**). In a drug repositioning screen of 1600 clinical compounds using HCT116 spheroid cultures, Senkowski et al. (2015) identified niclosamide, as well as nitazoxanide, as promising new anticancer agents. Wnt/β-catenin signaling is a principle driver of colorectal carcinomas (Ahmed et al. 2016) and niclosamide was found to downregulate Wnt signaling and elicit colorectal antitumor responses (Osada et al. 2011). To optimize this chemotype further, Mook et al. (2015) synthesized over 35 niclosamide analogs to understand its SAR for inhibition of Wnt and to identify compounds with improved systemic exposure. A comparison of the 304 niclosamide analogs we’ve characterized for TMEM16A activity (Supplementary Figure 4) found fifteen were identical to compounds they had characterized, were potent antagonists of *both* TMEM16A and Wnt signaling and had activity that correlated (Supplementary Figure 44). Niclosamide had an IC_50_ of 0.34 μM for inhibition of Wnt/β-catenin transcription using a TOPFlash reporter assay (Mook et al. 2015), which matches closely the IC_50_ of 0.13 μM we measured for inhibition of the TMEM16A, a molecular target also implicated in cancer. The molecular structure and activity of the fifteen niclosamide analogs is shown in **Figure 11D**. While niclosamide has a safe track record in man, compounds with a nitro group are often avoided due to potential toxicity concerns (Patterson and Wyllie 2014). Removal of the nitro group of niclosamide resulted in only minor reduction in TMEM16A activity (Cpd 11), while equivalent potency was observed when it was replaced with a chlorine group (Cpd 12). Several additional derivatives were identified having sub-μM potency on TMEM16A. While none showed improved activity compared to niclosamide, understanding what structural substitutions are tolerated is equally important to identify derivates with improved solubility, pharmacokinetics and stability *in vivo*.

## DISCUSSION

For the first time, we identify the approved drugs niclosamide and nitazoxanide as TMEM16A antagonists. With an IC_50_ of 132 nM in blocking the TMEM16A halide sensitive YFP response, niclosamide is one of the most potent antagonists described to date. To provide perspective on their activity, niclosamide and related compounds were compared to eight benchmark TMEM16A antagonists described earlier. The pharmacology of niclosamide and relateds in providing potent but partial block of the iodide eYFP response yet full block of the TMEM16A chloride current appears most similar to the benchmark antagonists 1PBC and NTTP. Thus, niclosamide and relateds may share their mechanism as pore blockers. While we describe the activity of seventeen niclosamide analogs, over 300 niclosamide analogs have been tested and the partial block of the iodide eYFP response is characteristic of this chemotype. Further investigation is needed to understand why these compounds show partial inhibition of the iodide/eYFP response, but our findings are consistent with those of Peters et al. (2015) who similarly found 1PBC and NTTP only partially inhibited the TMEM16A iodide/eYFP response but fully inhibited the calcium-activated chloride current (see Supplementary Figure S2 of their online article). Our detailed evaluation of niclosamide SAR and the background literature on this anthelmintic motivated us to also analyze the approved drug nitazoxanide for TMEM16A activity since nitazoxanide was synthesized in early 1970s using the niclosamide scaffold, had improved oral bioavailability and exhibited no significant impact on cytochrome P450 enzymes (White 2004, Anderson and Curran 2007, Rossignol 2014). Remarkably, nitazoxanide and its metabolic product tizoxanide also blocked TMEM16A with low single digit μM potency and showed a similar pharmacology to niclosamide in the eYFP and ephys assays. Upon further scrutiny of these compounds, we were surprised when we realized tizoxanide is a highly similar structural analog of NTTP, a known TMEM16A antagonist. We suggest this provides further evidence they may share this mechanism as pore blockers and propose they may interact with the same charged residues, Arg 511 and Lys 599, which Peters et al. (2015) find are important for NTTP and 1PBC inhibition of TMEM16A. These two basic residues along with Arg 531 which upon mutation to alanine alter anion selectivity of TMEM16A (Peters et al. 2015), have been confirmed recently in the cryoEM structures of mouse TMEM16A to reside near the external narrow neck of the channel pore where the positive charges facilitate anion entrance and permeability (Paulino, Kalienkova, et al. 2017, Paulino, Neldner, et al. 2017, Dang et al. 2017). To our knowledge the mechanism of action for the benchmark TMEM16A antagonists CaCCinh-A01, benzbromarone, dichlorophen and MONNA remains unknown, but because these compounds show a distinct pharmacology from niclosamide and relateds it is likely they have different site/s of action. Niflumic acid which has a pharmacology similar to niclosamide, 1PBC and NTTP was proposed over a decade earlier to be a pore blocker (Piper, Greenwood, and Large 2002) and to preferentially block larger versus smaller halides (Ni, Kuan, and Chen 2014). At this time, we’ve run only limited counterscreens for selectivity over other chloride channels, but promisingly find niclosamide and related compounds showed little impact on the CFTR chloride current, as did the pore blockers 1PBC and niflumic acid.

TMEM16A is expressed on airway smooth muscle cells which display the classical electrophysiology properties of CaCC (Gallos et al. 2013, Zhang et al. 2013), including a rapid calcium-dependent inactivation (Wang and Kotlikoff 1997) which we also observe after ionomycin-activation of cloned TMEM16A in HEK293 cells (**Figures 4G,H** and Supplementary Figure 25). We have confirmed TMEM16A expression using an RNAseq dataset (Aisenberg et al. 2016) of primary human ASM and bronchial epithelial (HBE) cells (**Figure 10**), detecting almost exclusively the ‘abc’ splice variant lacking segment ‘d’ in HBE cells as reported earlier (Caputo et al. 2008), but find ASM cells express a mixture of splice variants with variable amounts of segment ‘b’ and ‘d’ (Supplementary Figure 41). In this later case, this may include the ‘acd’ variant we used for QPatch electrophysiology studies, while in the former it would include the ‘abc’ variant we used in our high-throughput TMEM16A eYFP screen. While we’ve primarily used the ‘acd’ variant for patch clamp electrophysiology studies on antagonists, it’s worth noting the pore blocker 1PBC showed similar efficacy in blocking TMEM16A ‘acd’ and ‘abc’ Ca^2+^-activated Cl^-^ currents (**Figure 4C**, Supplementary Figures 11 and 24). Irrespective, since pro-contractiles such as histamine and methacholine activate calcium-activated chloride channels leading to membrane depolarization, calcium mobilization and contraction of smooth muscle cells, we advanced TMEM16A antagonists for direct interrogation of impacts on these processes.

Niclosamide and related compounds blocked histamine depolarization of ASM cells at sub-micromolar concentrations. These studies confirm earlier findings that TMEM16A antagonists hyperpolarize human ASM cells (Yim et al. 2013, Danielsson et al. 2014, Danielsson et al. 2015) and extend them to demonstrate antagonists also reverse the depolarization induced by contractants. To provide perspective on the activity of niclosamide and related compounds, we compared them to seven small molecule antagonists described earlier of distinct chemotypes (**Figure 2**). Consistent with TMEM16A’s role in controlling ASM excitation-contraction coupling, we find antagonists characteristically blocked pro-contractile depolarization and importantly provided robust bronchodilation of mouse tracheal rings and human small airways (**Figure 5**). In alignment with their enhanced potency on TMEM16A, niclosamide and relateds showed improved activity in relaxing airways. Our findings extend earlier studies demonstrating niflumic acid relaxes TEA or K+ gluconate pre-contracted guinea pig tracheal rings, rat small airways and human tracheal smooth muscle strips (Yim et al. 2013, Danielsson et al. 2014), benzbromarone and tannic acid suppress Substance P contraction of guinea pig tracheal rings (Gallos et al. 2013) and benzbromarone relaxes methacholine-contracted mouse peripheral airways or human airway smooth muscle strips pre-contracted with acetylcholine or LTD4 (Danielsson et al. 2015). Importantly, these findings and our results using carbachol and histamine indicate TMEM16A antagonists offer a new mechanism to block the multiple contractiles operating in disease. More recently using smooth-muscle-specific TMEM16A knockout mice, Wang et al. (2018) demonstrate agonist-induced inward currents and contractility of airway smooth muscle is profoundly reduced in the absence of TMEM16A, cementing its role as a key regulator of ASM excitation and contraction.

Asthma is a chronic inflammatory disorder characterized by airway hyperresponsiveness (AHR). Significantly, Zhang et al. (2013) find in mouse model of asthma, ovalbumin-sensitization upregulates TMEM16A expression in ASM cells and benzbromarone and niflumic acid prevented AHR and contractions evoked by methacholine. We also have devised schemes to evaluate the efficacy of TMEM16A antagonist under extreme conditions of contraction and inflammation that reduce β-agonist action (Lemoine and Overlack 1992, Shore et al. 1997). Remarkably, maximally contracted mouse airways were fully relaxed by the TMEM16A antagonists niclosamide, Cpd 4 and benzbromarone, while the β-agonist isoproterenol showed only partial bronchodilation which importantly converted to full bronchodilation with add-on of a TMEM16A antagonist (**Figure 8**). Severe asthma and COPD are also associated with poorly controlled inflammation, steroid insensitivity and reduced β-agonist responsiveness. Thus, there is a need for new bronchodilators with a distinct MOA that can operate amidst this inflammation. The cytokines IL-1β and TNFα were shown earlier to reduce β-agonist relaxation of airway smooth muscle (Shore et al. 1997, Moore et al. 2001). More recently bitter taste receptors were shown to provide a new mechanism to bronchodilate airways which resists IL-13 desensitization events limiting β-agonist action (Robinett et al. 2014). Using mouse tracheal rings treated with a more robust new cocktail of three cytokines, IL-1β, IL-13 and TNFα, we demonstrate β-agonist bronchodilation is profoundly reduced, yet TMEM16A antagonists remain remarkably effective in fully relaxing airways (**Figure 9**). Finally, because asthma treatment guidelines include β-agonists as a standard of care but allow for add-on therapies for poorly controlled disease, we investigated if TMEM16A antagonists can operate with β-agonists, including under situations of tachyphylaxis or use-dependent desensitization. Promisingly, airways exhibiting hyporesponsiveness after repeat β-agonist treatment were fully relaxed upon addition of the TMEM16A antagonist, Cpd 4.

Secretory epithelial cells, smooth muscle cells and sensory neurons express the calcium-activated chloride channel TMEM16A. In the bronchial epithelium, TMEM16A is strongly upregulated by Th2 cytokines which correlates with goblet cell hyperplasia and mucin hypersecretion, as well as, increased calcium-dependent chloride secretion (Scudieri et al. 2012, Huang et al. 2012, Caputo et al. 2008). We have confirmed these results finding TMEM16A is dramatically upregulated in the bronchial epithelium after IL-13 treatment and expressed on the apical surface of goblet cells, but also show that cynomolgus monkeys with allergic asthma have elevated TMEM16A staining within the submucosal glands (**Figure 10**) which are an underappreciated but major source for excessive mucin production in disease. Interestingly, like our findings with IL-13, Gorrieri et al. (2016) similarly find TMEM16A expression peaks quickly, 24 hours after IL-4 treatment, a time frame we find precedes maximum Muc5AC expression occurring 5-7 days later (**Figure 10**). Notably, using an IL-13 mouse model of asthma, Nakono et al. (2006) reported that niflumic acid inhibited goblet cell hyperplasia and airway hyperresponsiveness. Huang et al. (2012) extended these findings demonstrating TMEM16A is upregulated in the asthmatic bronchial epithelium and its inhibition with benzbromarone and dichlorophen blocked mucin secretion and ASM contraction. More recently, TMEM16A inhibition with antagonists or siRNA has been reported to reduce IL-13 induced Muc5AC production and goblet cell hyperplasia (Lin et al. 2015, Zhang et al. 2015, Qin et al. 2016), while TMEM16A overexpression had opposite effects in increasing Muc5AC expression (Lin et al. 2015, Qin et al. 2016). Thus, TMEM16A inhibitors may repress two of the key features leading to airway obstruction in severe asthma and COPD, bronchoconstriction and mucin hypersecretion.

In contrast for cystic fibrosis, activation of TMEM16A on the apical membrane of epithelial cells has been proposed as one strategy to bypass the CFTR chloride secretory defect to improve hydration and mucociliary clearance, but bronchoconstriction and pain have been pointed out as possible unwanted side effects (Sondo, Caci, and Galietta 2014, Liu et al. 2016). To evaluate the effects of direct channel activation, we employed the TMEM16A opener, Eact (Namkung et al. 2011), and investigated its effects on airway smooth muscle and the bronchial epithelium. Interestingly, Eact induced robust ASM cell depolarization and calcium flux like contractants, which was blocked by antagonists (Supplementary Figures 31 and 32). The average EC_50_ of 1.6 μM for Eact induced ASM depolarization, closely matched its EC_50_ of 3.0 μM in activating TMEM16A (Namkung et al. 2011). These findings are consistent with those from (Danielsson et al. 2015), who found Eact depolarized ASM cells like contractants, while antagonists hyperpolarized ASM cells. To investigate its effects on bronchial epithelial cells, we incubated mature bronchial epithelial air-liquid interface (ALI) cultures for 2-3 days with the Eact opener. Muc5AC and leukotriene (LTC4, LTB4) secretion from bronchial epithelial ALI cultures was significantly increased (2-3 fold and 6-12 fold, respectively, p<0.05) following treatment with the TMEM16A opener, Eact (Supplementary Figure 42). While these findings supported the concept that agonists may have reciprocal effects to antagonists, our enthusiasm with these results was tempered by recent findings Eact was also a direct activator of TrpV1 (Liu et al. 2016), which can physically associate with TMEM16A in sensory neurons to enhance pain sensation (Takayama et al. 2015). As TrpV1 is also expressed by bronchial epithelial and ASM cells (McGarvey et al. 2014, Yocum et al. 2017), it’s activation alone, or with TMEM16A, could also explain some of our findings with Eact. Interestingly, Benedetto et al. (2017) also recently report TMEM16A and CFTR physically interact and cross-regulate each other in differentiated epithelial cells. Tissue specific knockout of TMEM16A in mouse intestine and airways abolished not only Ca^2+^-activated Cl secretion, but also abrogated CFTR-mediated Cl secretion. Somewhat surprising, there was no overt phenotype despite the complete absence of chloride currents in knockout tissues and mucociliary clearance was not compromised, but unexpectedly enhanced in tracheas from knockout mice. While this may support airway Na^+^ transport is physiologically more relevant for hydration and TMEM16A knock-down may enhance mucociliary clearance, further studies are needed to understand TMEM16A’ s role under disease conditions and the effects of pharmacological agents in modulating goblet cell hyperplasia and mucin hypersecretion. Niclosamide probably inhibits TMEM16A in mucus producing cells which inhibits mucus production and / or secretion, however, there remains a need for TMEM16A openers to explore the consequence of direct channel activation.

Beyond respiratory disease, antagonists of TMEM16A have been proposed to have utility in treating a wide variety of other diseases including, pulmonary hypertension (Namkung et al. 2010, Forrest et al. 2012, Heinze et al. 2014), secretory diarrhea (Thiagarajah, Donowitz, and Verkman 2015, Ousingsawat et al. 2011), polycystic kidney disease (Buchholz et al. 2014), pain (Cho et al. 2012, Lee et al. 2014, Pineda-Farias et al. 2015) and cancer. TMEM16A has been implicated in numerous cancers, including head and neck, esophageal, gastric, colorectal, lung, breast, prostate, pancreatic and glioma (reviewed recently by (Wang et al. 2017)). TMEM16A overexpression contributes directly to tumorigenesis and cancer progression and its knockdown with shRNA or small molecule antagonists ameliorates disease (Britschgi et al. 2013, Liu et al. 2012, Duvvuri et al. 2012, Wang et al. 2017). It associates with EGFR to regulate cancer cell proliferation and activates several pathways important for tumor growth (Britschgi et al. 2013, Bill et al. 2015). Interestingly, drug repurposing efforts have similarly found the anthelmintics niclosamide and nitazoxanide hold great promise for the treatment of cancer, as well as numerous other disorders, including hypertension, secretory diarrhea, and as a broad-spectrum anti-infective agent (Chen et al. 2017, Rossignol 2014, Li et al. 2014, Rossignol et al. 2006). Underlying mechanisms proposed for its action have included uncoupling of oxidative phosphorylation, modulation of Wnt/β-catenin, mTORC1, STAT3, NF-κB and Notch signaling, but identifying a unifying mechanism for niclosamide’s action has remained elusive (Chen et al. 2017). We’d suggest antagonism of TMEM16A should now be considered as a possible contributing factor. This is especially so for secretory diarrhea and cancer where the role of TMEM16A has been well established. Activation or upregulation of the calcium-activated chloride channel TMEM16A, an upstream target on the cell surface, can initiate a variety downstream signaling pathways affecting various pathophysiological processes (e.g. as shown for Ras-Raf-MEK-ERK in cancer (Duvvuri et al. 2012). Receptor-mediated increases intracellular calcium have also been found to be Cl^-^ dependent and TMEM16A antagonists have been found to modulate calcium homeostasis (Cabrita et al. 2017, Danielsson et al. 2015). Channels can also associate with other cell surface molecules to form large macromolecular signaling complexes. Indeed, TMEM16A has been shown to interact with the ezrin-radixin-moesin network (Perez-Cornejo et al. 2012) and with TrpV1 (Takayama et al. 2015), TrpC6 (Wang et al. 2016) and the IP3 receptor (Jin et al. 2013) for coupled GPCR and ion channel signaling. Interestingly, the key signaling molecule β-catenin involved in numerous cancers was also found recently to associate with the ion channels KCNQ1 (Rapetti-Mauss et al. 2017) and BKCa (Bian et al. 2011) to modulate Wnt signaling.

It is important to mention some limitations of our research. Our work was focused in broadly evaluating many compounds to develop a structure-activity relationship and advance a lead, as opposed to a more narrow but deep understanding of the pharmacology and mechanism of any single compound. Future studies to better understand niclosamide and nitazoxanide’s impact on permeation of different anions, their activity on native chloride channels, specificity and mechanism of block, and *in vivo* activity are warranted. This is especially so given the wide interest in these compounds from a drug repositioning standpoint and our new discovery they are TMEM16A antagonists. The bioavailability and suitability of these compounds for respiratory disease also needs to be explored in greater detail. While the short-acting β-agonists (SABA) salbutamol and isoproterenol are hydrophilic (XLogP3 of 0.31 and −3.49, respectively) and act quickly (<7 min) to open airways which is necessary for a rescue treatment, the TMEM16A antagonists niclosamide, Cpd 4 and benzbromarone are quite lipophilic (XLogP3 of 5.32, 4.63, and 5.66, respectively), have a slow onset of action (17-60 min) but may show sustained effects in bronchodilating mouse airways. The long-acting β-agonist (LABA) salmeterol is also quite lipophilic (XLogP3 of 3.90) and has been similarly observed to have a slow onset of action (90 min) but sustained effects in relaxing airways (Stocks et al. 2014). Such compounds, however, can be challenging from a development standpoint as they tend to have poor solubility, but they have ideal characteristics as an inhalation maintenance therapy since they offer long-lasting effects and are retained in the lung which is a safety advantage. Whether niclosamide and related compounds have similar advantages should be examined further, including if they share the long duration of action of the LABA salmeterol, which is believed principally to be the result of a physicochemical interaction of the lipophilic compound with membrane lipid bilayers acting as a depot to allow drug rebinding, the so-called ‘plasmalemma diffusion microkinetic model’ (Anderson, Linden, and Rabe 1994). While the majority of TMEM16A antagonists were lipophilic, nitazoxanide and tizoxanide with XLogP3 values of 2.04 and 3.15, respectively, were unique in being somewhat more hydrophilic. These compounds merit further explorations as they may offer some development advantages. Additional research to better understand the molecular interaction of antagonists with TMEM16A is also warranted to improve compound activity and clarify their mechanism in blocking the channel.

TMEM16A antagonists offer an exciting new strategy to treat a wide variety of disorders. We demonstrate antagonists block airway smooth muscle depolarization and contraction, offering a promising new mechanism to bronchodilate airways that resists use- and inflammatory-desensitization pathways limiting β-agonist efficacy in severe asthma and COPD. We identify the approved drugs niclosamide, nitazoxanide and related compounds as some of the most potent TMEM16A antagonist described to date and provide a molecular target to compounds of high interest from drug repurposing efforts. Further studies will be needed to assess their efficacy in these new disease indications, but niclosamide and nitazoxanide’s safe track record in man position these chemical series as a good starting point for additional studies.

## ADDITIONAL INFORMATION

### Author Contributions

A.H., K.L. and K.W. developed and conducted the high-throughput screen and performed follow-up experiments to identify validated hits. P.W. and D.P. performed medchem support in determining compound potency in eYFP assay. B.L., K.H., J.M. and J.KS. developed TMEM16A electrophysiology assays, performed experiments and/or analyzed data. K.G. and X.X. developed and analyzed compounds for effects on ASM membrane potential or calcium flux. G.Y. characterized compounds for effects on HBE ALI cultures. K.M., R.E., D.M. and A.L. developed and characterized compounds for bronchodilation of mouse, cynomolgus monkey or human tissue. K.M. ran routine bronchodilation studies to support medchem and performed studies evaluating compound ability to resist use- and inflammatory-desensitization. E.T., K.H., S.E., O.H., K.M., B.B., J.W., and M.F. conducted experiments evaluating TMEM16A expression and splice variants. J.K.S., D.S., M.C., T.B., A.B., J.J., J.P., M.W. and K.K. were involved in conception, data analysis or enablement. J.C. and L.L. synthesized compounds and analyzed data. J.K.S. initiated the project, supervised the program and wrote the first draft of the manuscript; while A.H. and K.K. made critical edits on the final draft. All authors discussed and commented on the manuscript.

### Author ORCIDs

Andreas Hochheimer, iD https://orcid.org/0000-0002-1572-9334 Deanna Mohn, https://orcid.org/0000-0001-5748-7591 John K. Sullivan, https://orcid.org/0000-0001-9840-852X

### Present Address

Katja Labitzke, Clariant Produkte (Deutschland) GmbH, Planegg, Germany.

Longbin Liu, CHDI Management, CHDI Foundation, Los Angeles, CA, USA.

Anh Leith, Dept. Genome Sciences, University of Washington, Seattle, WA, USA.

Esther Trueblood, Seattle Genetics, Bothell, WA, USA.

Teresa L. Born, Sartorius Stedim BioOutsource, Cambridge, MA, USA.

Alison Budelsky, Immunology Research, Lilly Research Laboratories, San Diego, CA, USA.

Dirk Smith, Insight Bioconsulting, Bainbridge Island, WA, USA

Kerstin Weikl, Assay.Works GmbH, Regensburg, Germany.

Andreas Hochheimer, ISAR Bioscience GmbH, Planegg, Germany.

John K. Sullivan, PolestarBio LLC, Newbury Park, CA, USA.

### Ethics

Animal experimentation and tissue: All animal procedures and tissue collections were conducted in an Association for Assessment and Accreditation of Laboratory Animal Care accredited facility in accordance with the requirements and guidelines of the US National Research Council and in compliance with the protocols approved by the Institutional Animal Care and Use Committee of the following institutions: Amgen Inc., Thousand Oaks, CA; Charles River, Shrewsbury, MA.

All human lung specimens were collected under Institutional Review Board approval with appropriate informed consent. In all cases, materials obtained were surplus to standard clinical practice. Patient identity and PHI/identifying information were redacted from tissues and clinical data.

## ACKNOWLEDGMENTS

The authors wish to thank the Amgen Protein Technology group for TMEM16A expression constructs, the scientific team at Sophion and ChanTest (CRL) for help in developing the QPatch and IonWorks Barracuda electrophysiology assays or counterscreens against CFTR, scientists at Biopta (ReproCELL) for bronchodilation studies, and past members of the Airway Smooth Muscle Working Group, Asthma, COPD and IPF Teams at Amgen who contributed to building a foundation in the respiratory efforts which enabled these studies, including Heather Thomas, Ken Schooley, Ryan Brown, Deb Hopkins, Anna Pirrone, Leanne Peiser, Cindy Willis, Daniela Metz and Heather Arnett.

## SUPPLEMENTARY MATERIAL

The Supplementary Material for this article can be found online at:

